# Comparative Parallel Multi-Omics Analysis During the Induction of Pluripotent and Trophectoderm States

**DOI:** 10.1101/2020.09.27.315259

**Authors:** Mohammad Jaber, Ahmed Radwan, Netanel Loyfer, Mufeed Abdeen, Shulamit Sebban, Thorsten Kolb, Marc Zapatka, Kirill Makedonski, Aurelie Ernst, Tommy Kaplan, Yosef Buganim

**Author notes:** Correspondence should be addressed to Y.B. or T.K. These authors contributed equally to this work (Mohammad Jaber, Ahmed Radwan, Netanel Loyfer). Lead contact: Yossi Buganim.

## Abstract

Following fertilization, totipotent cells divide to generate two compartments in the early embryo: the inner cell mass (ICM) and trophectoderm (TE). It is only at the 32-64 -cell stage when a clear segregation between the two cell-types is observed, suggesting a ‘T’-shaped model of specification. Here, we examine whether the acquisition of these two states *in vitro* by nuclear reprogramming share similar dynamics/trajectories. We conducted a comparative parallel multi-omics analysis on cells undergoing reprogramming to Induced pluripotent stem cells (iPSCs) and induced trophoblast stem cells (TSCs), and examined their transcriptome, methylome, chromatin accessibility and activity and genomic stability. Our analysis revealed that cells undergoing reprogramming to pluripotency and TSC state exhibit specific trajectories from the onset of the process, suggesting ‘V’-shaped model. Using these analyses, not only we could describe in detail the various trajectories toward the two states, we also identified previously unknown stage-specific reprogramming markers as well as markers for faithful reprogramming and reprogramming blockers. Finally, we show that while the acquisition of the TSC state involves the silencing of embryonic programs by DNA methylation, during the acquisition of pluripotency these specific regions are initially open but then retain inactive by the elimination of the histone mark, H3K27ac.

## Cellular Specification during Early Embryogenesis

Fertilization of an oocyte by a sperm initiates robust epigenetic reprogramming of the DNA content within the newly formed cell, resulting in a totipotent zygote that holds the potential to produce all embryonic and extra-embryonic tissues^1^. Several divisions later, an early blastocyst is formed, containing two compartments that are more committed: an inner cell mass (ICM) which contains pluripotent cells (epiblast (Epi)) that will form the embryo proper, and an outer layer of trophectoderm (TE) cells, which will give rise to components of extra-embryonic tissues such as the placenta^2–4^.

Exactly how the specification between the ICM and TE cells is made during embryogenesis is not fully understood, although several models have been suggested^2–4^. Recently, the transcriptional trajectory from zygote to blastocyst has been described using single-cell transcriptomic data^5–11^. Interestingly, while clear transcriptional changes are found between the different stages (i.e. zygote, 2-cell stage, 4-cell stage, 8-cell stage, morula and blastocyst), the transcriptional heterogeneity within the cells of each group before blastocyst formation is relatively mild. Although some genes, like *Sox21*, were shown to exhibit transcriptional heterogeneity even within the 4-cell stage^6^, the overall transcriptome is relatively similar between the two groups of cells. This suggests a ‘T’-shaped model, where cells at each stage undergo relatively similar transcriptional changes before segregation, and separate into two distinct cells types, the ICM and TE, only at the morula/early blastocyst stage. This notion is supported by multiple evidence; first, each of the 2-8 first cells of the embryo can give rise to both TE and ICM. Second, cells of the outer layer of the morula stage have been observed to migrate into the inner layer and become pluripotent cells. This suggests a dynamic chromatin landscape and transcriptome that are relatively analogous between the cells before the final specification^3, 4^.

## Somatic Epigenetic Reprogramming to Pluripotency and Trophoblast Stem Cell State

Epigenetic reprogramming of a somatic nucleus to pluripotency or to a TE state has been achieved *in vitro* by somatic cell nuclear transfer (SCNT^12^) or by forced expression of a defined number of transcription factors^13–19^. While ectopic expression of Oct4, Sox2, Klf4 and Myc (OSKM) induces the formation of pluripotent stem cells (PSC, the *in vitro* counterpart of the ICM-Epi)^17^, we and others have shown that ectopic expression of Gata3, Eomes, Tfap2c and Myc (GETM, or Ets2 instead of Myc) induces the formation of trophoblast stem cells (TSC, the *in vitro* counterpart of the TE)^13, 16^. Importantly, in both reprogramming systems, the resulting cells are equivalent to their *in vitro* blastocyst-derived counterparts in their transcriptome, epigenome and function^13, 15–17, 20^.

However, while nuclear reprogramming to pluripotency and TE state during fertilization or in SCNT occurs within 2-3 days^21^, nuclear reprogramming by defined transcription factors is a long and inefficient process^13, 15–17^. Intrigued by these fundamental differences, scientists have devoted the last decade to monitoring and describing the various mechanisms, stages and pathways that underlie somatic cells undergoing reprogramming to pluripotency^14^. These major efforts have revealed key aspects in nuclear reprogramming which also explain, at least partially, the low efficiency of the process and describe in detail the trajectories somatic cells undergo in their way to become iPSCs.

However, while some properties of the reprogramming process to pluripotency have been illuminated, the characterization of the reprogramming process to the TSC state, or more intriguing, concomitant comparative multi-omics analysis of cells undergoing reprogramming to the TSC and pluripotent states has never been performed.

Here, we describe the trajectories and key elements that regulate and characterize cells undergoing reprogramming to iTSCs and iPSCs in depth. We show that in contrast to early embryonic cells, fibroblasts transduced with GETM or with OSKM mostly follow a ‘V’-shaped model where cells acquire, from the onset of the reprogramming process, a unique and specific chromatin and transcriptional programs that are mostly mutually exclusive and important for the induction of each state. Surprisingly, this ‘V’-shaped behavior was also evident at the methylation levels, where correlation with transcription is relatively low. Using single-cell analysis, we revealed unique and previously unknown markers for each reprogramming system. Moreover, chromatin accessibility and activity analyses identified many global reprogramming blockers such as Usf1/Usf2, Nrf2 and MafK along with other oxidative stress response genes that significantly hinder both reprogramming systems but with a different dynamic.

Lastly, by integrating all the data together we illuminated key aspects that characterize each fate. Remarkably, by comparing both systems we show that from the onset of the reprogramming process OSKM define regions that are important for the development of the heart and brain, two most essential organs for the developing embryo. Moreover, we show that while GETM shut off the embryonic program by DNA methylation, OSKM open these regions but retain them as inactive by eliminating the histone mark H3K27ac. In conclusion, besides providing the first multi-layer characterization of cells undergoing reprogramming to the TSC state, our approach of conducting concomitant and comparative multi-omics analysis of cells acquiring both pluripotency and TSC state allowed us to identify previously unknown properties for OSKM reprogramming as well.

## Results

### The Trajectory from the Zygote to the Blastocyst Stage is following a ‘T’-Shaped behavior

The trajectory from the zygote to the blastocyst stage (**Fig. 1A**) has recently been described by several studies using single-cell transcriptomic data^5–11^. Using principal component analysis (PCA), Deng et al. suggested that the trajectory from zygote to blastocyst follows a ‘U’ shape^5^, in which PC1 separates between the zygote/early 2-cell stage and blastocyst and PC2 separates between the other stages, namely the 2-16-cell stage and the zygote/blastocyst (**Fig. 1B**). However, since the zygote and the 2-cell stage are considered totipotent and thus harbor a unique transcriptome, we reanalyzed the data by excluding these two stages. Interestingly, the re-analyzed PCA revealed a clear ‘T’-like shape where PC1 separates between the 4-cell stage and the blastocyst and PC2 between the ICM and TE (**Fig. 1C** **and Extended Data Fig. 1A**). More importantly, while both analyses (**Fig. 1B-C**) suggest that a clear transcriptional shift between different stages occurs during early embryogenesis, in both analyses the heterogeneity within each group was mild, indicating that the cells undergo relatively similar changes during embryogenesis and before specification. A ‘T’-shaped behavior of early embryonic cells was similarly observed in the datasets of Guo et al.^7^, strengthening the notion that only at the morula/early blastocyst stage, a clear transcriptional segregation between cells of the same developmental stage can be witnessed (**Fig. 1D** **and Extended Data Fig. 1B**).

**Fig. 1.**
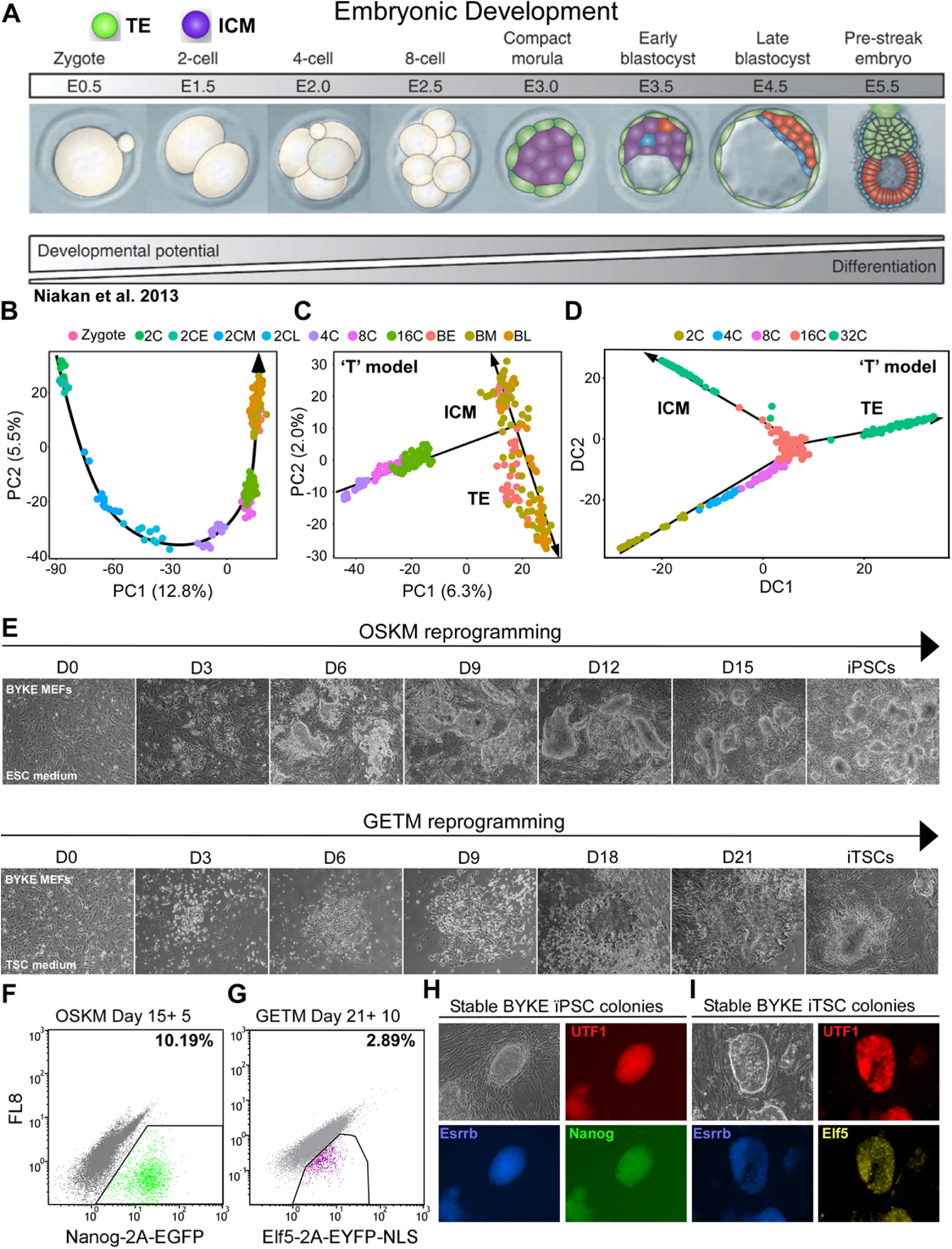
Establishment of the pluripotent and trophectoderm states in the embryo and during somatic nuclear reprogramming. **(A)** An illustration of the various early embryonic stages and forming cell types during embryogenesis (adapted from^108^). Inner cell mass (ICM, purple) and trophectoderm (TE, green) are the first compartments to show a clear transcriptional specification. **(B + C)** Single-cell RNA sequencing data obtained from different stages of developing embryo^5^ demonstrating the trajectory from zygote to blastocyst. PCA graphs showing gene expression profiles among 252 single cells projected onto the first two principal components. The trajectory from the zygote to the blastocyst is following a U-like shape (B) while the exclusion of totipotent cells (zygote and 2-cell stage (2C)) allows the visualization of a T-like shape progression segregating the TE from the ICM. **(D)** Single-cell RNA sequencing data obtained from different stages of developing embryo^7^ demonstrating the trajectory from the zygote to the blastocyst stage. Diffusion map was constructed by MERLoT package using 48 genes in 433 individual cells obtained from 2C through blastocyst. Again a clear T-like shape progression is noted separating the ICM from the TE. **(E)** Representative bright field images showing cell morphology and cell density during OSKM reprogramming toward iPSC formation (top) and during GETM reprogramming toward iTSC generation (bottom). **(F)** FACS analysis for Nanog-2A-EGFP reporter on BYKE MEFs undergoing reprogramming for 15 days with OSKM factors followed by 5 days of dox removal. **(G)** FACS analysis for Elf5-2A-EYFP-NLS reporter on BYKE MEFs undergoing reprogramming for 21 days with GETM factors followed by 10 days of dox removal. **(H)** Bright field and fluorescence images of a stable BYKE iPSC colony demonstrating the activation of the 3 pluripotent reporters (Utf1-2A-tdTomato/Esrrb-2A-TagBFP/Nanog-2A-EGFP). **(I)** Bright field and fluorescence images of a stable BYKE iTSC colony demonstrating the activation of the 3 TSC reporters (Utf1-2A-tdTomato/Esrrb-2A-TagBFP/Elf5-2A-EYFP-NLS).

We sought to determine whether this ‘T’-like behavior also characterizes somatic cells undergoing reprogramming to pluripotency and TSC state. In general, the reprogramming process of fibroblasts is characterized by multiple steps: (1) loss of the somatic cell identity, (2) rapid proliferation, (3) mesenchymal to epithelial transition (MET), (4) metabolic shift, (5) stochastic gene expression, (6) epigenetic switch, (7) silencing of the exogenous factors by methylation and finally (8) the stabilization of the core cellular circuitry^14, 32^. We proposed three possible models, ‘T’, ‘Y’ and ‘V’, which may represent the trajectory of fibroblasts undergoing reprogramming into iPSCs and iTSCs (**Extended Data Fig. S1C**). The ‘T’-shaped model predicts that fibroblasts undergoing reprogramming into iPSCs or iTSCs will undergo comparable transcriptional and epigenetic changes during the conversion and that a separation between the two cell types will occur only at the end of the reprogramming process, similar to the cells of the early embryo. The ‘Y’-shaped model predicts that only genes and regulatory elements that are responsible for early and general processes like loss of somatic cell identity, proliferation and MET will be shared between the two systems, after which each process will take a different path toward its own fate. The ‘V’-shaped model predicts that although some early processes are the same between the two systems, each process will mostly employ different set of genes and regulatory elements to achieve its own unique fate.

To understand which of the three proposed models most accurately represents the reprogramming process toward the two states, we performed a parallel comparative multi-omics analysis on fibroblasts undergoing reprogramming to iPSCs by OSKM or to iTSCs by GETM factors simultaneously (**Fig. 1E**). We utilized our previously developed system for distinguishing between pluripotency and TSC reprogramming, BYKE mouse embryonic fibroblasts (MEFs), which contain 4 unique knock-in fluorescent reporters: (1) Nanog-2A-EGFP, a cytoplasmic reporter that marks specifically pluripotent cells, (2) Elf5-EYFP-NLS, a nuclear reporter that marks specifically TSCs and (3) Utf1-2A-tdTomato and (4) Esrrb-2A-TagBFP, cytoplasmic reporters that mark both cell types (**Fig. 1F-I**, ^42^). We then measured the transcriptome (bulk RNA-seq and single-cell RNA-seq (SC-RNA-seq)), methylome (reduced representative bisulfite sequencing, RRBS), chromatin accessibility (ATAC-seq), chromatin activity (ChIP-seq for H3K4me2 and H3K27ac) and genomic stability (CNVs) at different time points along the reprogramming processes and compared between the two systems (**Extended Data Fig. 1D**).

### Bulk Transcriptomic Analysis Depicts Unique Transcriptional Profiles for Each Reprogramming system

The transcriptional landscape of reprogrammable cells is the easiest and most robust examination to compare two parallel reprogramming systems (i.e. iPSC vs iTSC reprogramming). To explore the global transcriptomic changes of cells undergoing reprogramming to the pluripotent and TSC states, we reprogrammed BYKE MEFs into iPSCs with OSKM and into iTSCs with GETM and collected cell pellets in duplicates every three days along the reprogramming process (**Extended Data Fig. 1D**). The parental BYKE MEFs, the final cells (i.e. iTSCs and iPSCs) and their blastocyst-derived control counterparts (i.e. bdTSCs and ESCs) were analyzed as well.

We used full-transcript RNA-seq to estimate expression levels and transcribed isoforms, with a total depth of 20M reads per replicate. Initially we conducted principal component analysis (PCA) to observe the trajectory cells undergo during the reprogramming process to iTSCs and iPSCs (**Fig. 2A-F**). In addition, we extracted the gene loadings associated with the first two principal components in each PCA plot to reveal those genes that drive the distinction between the different stages/steps of each reprogramming process and between the two reprogramming systems (**Extended Data Fig. 1E-J**). Notably, cells undergoing reprogramming to a TSC or pluripotent state exhibited a very different transcriptional landscape, as analyzing both processes together in the same PCA plot, generated a ‘V’-like shape, starting from the beginning of the process **(Figs. 2C and 2F and Supplementary Figs. 1G and 1J**). This suggests that major transcriptional changes separate the two systems.

**Fig. 2.**
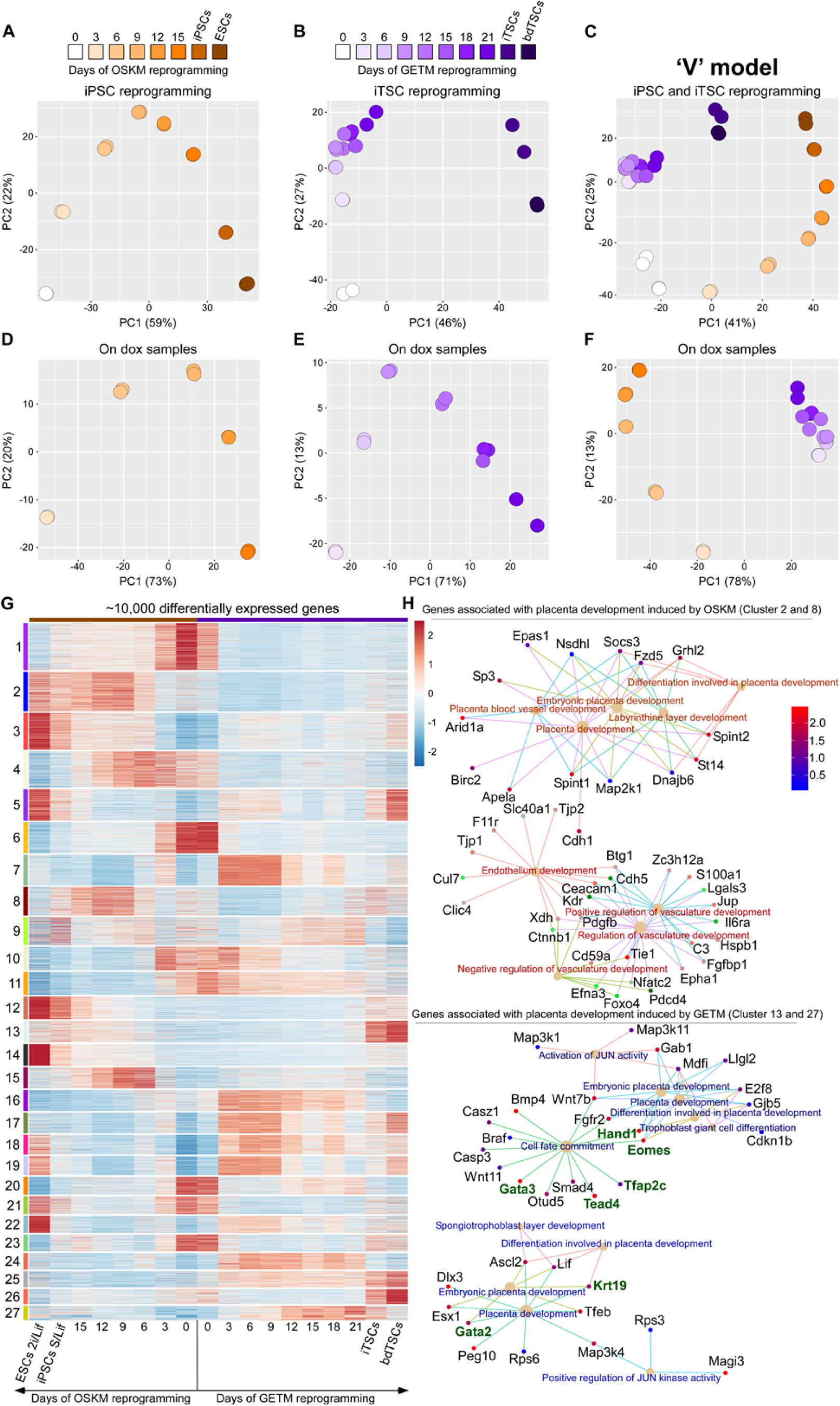
Bulk RNA-seq analysis on cells undergoing reprogramming to iPSCs and iTSCs suggests a unique transcriptional profile for each reprogramming system. **(A-C)** PCA plots describing the trajectory MEFs undergo during reprogramming to either iPSCs (A), iTSCs (B) or both (C) as assessed by gene expression profiles of bulk RNA-seq data projected onto the first two principal components. **(D-F)** same as in (A-C) but here only reprogrammable cells (cells on dox) are plotted. **(G)** Heatmap showing gene expression levels detected by bulk RNA-seq of 10,000 most variable genes among ESCs, bdTSCs, MEFs and cells during reprogramming. Unsupervised hierarchical clustering was performed and adaptive branch pruning was used to identify 27 prominent clusters. **(H)** Gene-concept network of GO terms associated with placenta development induced by OSKM (upper panel) or by GETM (lower panel) reprogramming. Key regulators of the trophoblast stem cell state such as Gata3, Gata2, Tfap2c, Tead4 and Eomes (marked by green) are only specific to the GETM reprogramming.

Interestingly, while the reprogramming process toward iPSCs showed constant and gradual transcriptional changes along the process until the stabilization of the final cells (**Figs. 2A and 2D and Supplementary Figs. 1E and 1H**), the reprogramming process toward iTSCs showed two main waves of transcriptional change where the first wave occurred very early (i.e. already at day 3, PC2, **Figs. 2B and Extended Data Fig. 1F**), followed by subtle changes in transcription until day 21 (**Fig. 2E** **and Extended Data Fig. 1I**) and a second wave which is initiated following transgene expression removal and is important for the activation of the core TSC circuitry (i.e. PC1, **Fig. 2B** **and Extended Data Fig. 1F**).

We believe that these differences between OSKM and GETM reprogramming are partially due to the nature of each reprogramming process. In our experience, while iPSC colonies may be stabilized during the reprogramming process and in the presence of transgenes, iTSC colonies cannot. Only when transgenes expression is shut off (i.e. removal of dox) stable iTSC colonies emerge.

Next, we took the ∼10,000 most differentially expressed genes among all samples and clustered them together, yielding 27 unique clusters (**Fig. 2G** **and Supplementary Table 1**). Clusters 1, 4, 6, 10, 11, 20 and 23 contain MEF-specific genes (i.e. gene ontology (GO) terms of connective tissue development, actin filament organization, extracellular matrix organization and angiogenesis) that are downregulated during GETM and OSKM reprogramming but with unique dynamics for each cluster and system (**Supplementary Table 1**). Clusters 7, 16, 17, 18, 19, 24, 25 and 27 are specific to the TSC reprogramming process. Interestingly, most of these clusters involve genes which are important to metabolism and cell cycle regulation (e.g. GO terms of ribonucleoprotein complex biogenesis, tRNA/rRNA/ncRNA metabolic processes and regulation of mitotic cell cycle, **Supplementary Table 1**). Clusters 2, 8, 12 and 15 are specific to iPSC reprogramming and contain genes that participate in cell junction organization, Ras protein signal transduction, regulation of vasculature development, histone modification and Wnt signaling (**Supplementary Table 1**). Cluster 3, 5 and 9 are shared between the two processes and compose of genes that regulate cell cycle, DNA repair and Wnt signaling pathway (**Supplementary Table 1**). We noticed that most genes behaved differently between the two reprogramming systems, even in early and shared dynamics such as proliferation, chromatin remodeling and mesenchymal to epithelial transition (MET, **Extended Data Fig. 2A-D**). While key mesenchymal genes and regulators of epithelial to mesenchymal transition (EMT) are downregulated in both systems as expected (**Extended Data Fig. 2C**, bottom of the heatmap), indicating loss of fibroblastic identity, particular mesenchymal and MET-specific genes are uniquely expressed in each reprogramming system (**Extended Data Fig. 2C-D**).

Another example for an important difference which can be observed already in early stages of reprogramming between the two systems is a metabolic shift which occurs in two waves (i.e. day 3-9 and days 12-21) in iTSC reprogramming. This shift, which plays a role in translation regulation activity and RNA processing, is completely absent in iPSC reprogramming (**Extended Data Fig. 2E-G and Supplementary Table 1**). Moreover, even when similar GO terms were annotated between the two systems, each reprogramming system utilized a different set of genes to execute the process. Figure 2H shows an example where both iTSC and iPSC reprogramming systems activated placenta/trophoblast-specific genes with ‘placenta development’ GO annotation, though while OSKM activated trophoblast differentiation genes, GETM activated trophoblast stem cell genes (marked by green, **Fig. 2H** **and Supplementary Table 1**). In conclusion, bulk transcriptomic analyses suggest a ‘V’-shaped behavior where GETM and OSKM operate distinctively to reprogram the somatic nucleus and already at the onset of the process take completely different transcriptomic routes, with minimal overlapping genes and signaling pathways.

### Single-Cell Transcriptomic Analysis Suggests No Overlap between GETM and OSKM Reprogramming

Bulk transcriptional analysis is a powerful tool to describe the global transcriptional changes that occur in most cells during the reprogramming process. However, it is lacking the sensitivity to identify the small fraction of cells that are destined to become iPSCs or iTSCs (typically only 1-5% of the cells for iTSC reprogramming and 10-20% for iPSC reprogramming acquire the final fully reprogrammed state^13, 42^, **Fig. 1F-G**). Moreover, due to population averaging, one cannot detect a small group of cells that might harbor a transcriptional profile that is similar between the two reprogramming systems, suggesting a ‘T’ or ‘Y’ behavior. To overcome this limitation, we conducted single-cell analysis on GETM and OSKM reprogrammable cells. While multiple studies employed the single-cell RNA-seq (SC-RNA-seq) technology to probe the transcriptome of individual cells undergoing reprogramming to pluripotency^34, 35, 43, 44^, similar characterization of the reprogramming process toward the TSC state or a comparative analysis between the two processes has never been done before.

To evaluate the transcriptomes of individual cells undergoing reprogramming into iPSCs and iTSCs, we exploited the 10X Genomics platform and profiled the transcriptome of ∼16,000 single cells at two time points (days 6 and 12) along the reprogramming process toward iPSCs and iTSCs. We chose these time points as they represent the stochastic gene expression phase that occurs in the two reprogramming systems and thus are expected to show the highest variation between individual cells. UMAP analysis for both days, 6 and 12, demonstrated two distinct clusters of cells; one for GETM and one for OSKM reprogramming (**Fig. 3A-B**). No overlapping cells were found between the two systems, indicating that each process acquired a completely different transcriptional profile, again suggesting a ‘V’-shaped model. Using EnrichR mouse gene atlas we identified the different transcriptional fates reprogrammable cells undergo during each reprogramming process. As the reprogramming process to iTSCs is much less efficient compared to OSKM reprogramming, we mostly identified non-reprogrammable MEFs (p≤8.9e-58) in the GETM reprogramming process with a very small fraction of cells with MEF-like transcriptional profile in OSKM reprogramming (p<4.1e-7, cluster 1 in **Extended Data Fig. 3A-B**). Interestingly, both processes contained cells with a transcriptional profile partially similar to that of the placenta (Cluster 3 for GETM and cluster 14 for OSKM, **Fig. 3A-B** and **Extended Data Fig. 3A**). This is in agreement with the bulk transcriptomic analysis that showed gene ontology of placenta development with unique gene set for each system (**Fig. 2H**). Other than MEFs and placental cells, different group of reprogrammable cells activated genes that are enriched in epidermis, dorsal root ganglia, mast cells and bone marrow in OSKM reprogramming and umbilical cord, bladder and NK cells in GETM reprogramming (**Fig. 3A-B** and **Extended Data Fig. 3A-B**). Epidermis, placenta and neuronal fates have previously been observed in OSKM reprogramming, strengthening our findings^43^. We next identified known and unknown stage-specific markers for each reprogramming process (**Fig. 3C-F**). For iTSC reprogramming, non-infected MEFs or refractory MEFs were identified using the known mesenchymal markers Thy1, Col1a2 and Postn. Bgn, Tagln and Scmh1 mark both MEFs and cells that have succeeded to initiate the reprogramming process. Dusp9, Dlk1, Cdca3 and Bex4 mark most reprogrammable cells that are in the midst of the reprogramming process prior to any fate decision, and Cbs, Wnt6, Pgf, Peg10, Cd82, Adssl1, Stard10 and Ppp2r2c mark cells that are either differentiated trophoblasts (Cluster #3) or those that are probably destined to become iTSCs, based on their unique intermediate stemness gene signature (part of cluster 5 and the junction between 5 and 3, **Fig. 3C-D** and **Extended Data Fig. 3C**).

**Fig. 3.**
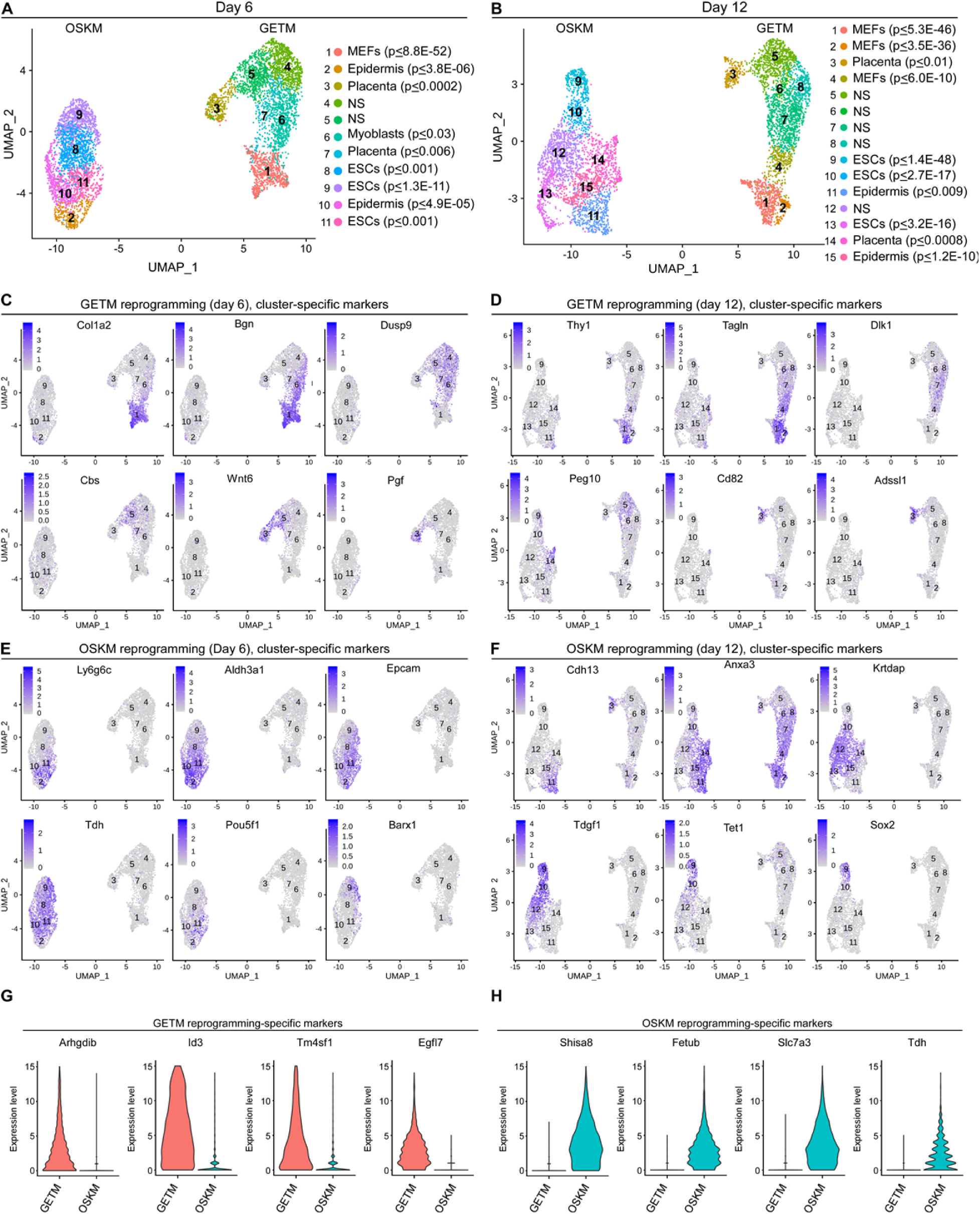
Single-cell RNA-seq analysis separates OSKM from GETM reprogramming. **(A)** Uniform Manifold Approximation and Projection (UMAP) visualization analysis of 5,899 single cells at day 6 of both OSKM and GETM reprogramming. Each point represents a single cell and each color represents a unique community among the population. 1000 marker genes were used to characterize all subpopulation and using the Mouse Gene Atlas (“ARCHS4”: https://amp.pharm.mssm.edu/archs4/; “MGA”: Mouse Gene Atlas), the closest significant cell type was assigned to each subpopulation. **(B)** UMAP visualization analysis of 5,752 single cells at day 12 of both OSKM and GETM reprogramming. 1217 marker genes were used to characterize all subpopulation and using the Mouse Gene Atlas, the closest significant cell type was assigned to each subpopulation. **(C + D)** Expression level of selected cluster-specific markers of GETM reprogramming at day 6 and day 12, respectively. The expression level of the specified markers is visualized by a range of intensities of a purple color. **(E + F**) Expression level of selected cluster-specific markers of OSKM reprogramming at day 6 and day 12, respectively. **(G + H)** Violin plots summarizing single-cell expression level of GETM-specific markers (G) and OSKM-specific markers (H) in GETM and OSKM reprogramming processes.

For OSKM reprogramming, Ly6g6c, Cd13, Krt16, Ecam1 and Cd34 represent cells that underwent MET and have acquired a partial epidermal fate. Krtdap, Epcam, and Tdh are markers for most OSKM reprogrammable cells but cannot predict successfully which cells are destined to become iPSCs. Interestingly, while Anxa3 marks all the cells that took a failed route toward pluripotency, high levels of Tdgf1 and Cenpf represent a successful trajectory to reprogramming based on gene expression (**Fig. 3E-F** and **Extended Data Fig. 3D**). As we and others have previously noted^14, 23, 45^, while Sox2, Dppa5a and to a lesser extent Tet1 stringently mark fully reprogrammed cells, Oct4 fails to do so (**Fig. 3E-F** and **Extended Data Fig. 3D**).

We next searched for genes that can robustly distinguish between GETM-reprogrammable cells and OSKM-reprogrammable cells. We identified four unique and specific markers for each system. While Arhgdib, Id3, Tm4sf1 and Egfl7 mark specifically most GETM-reprogrammable cells, Shisa8, Fetub, Slc7a3 and Tdh are uniquely expressed in almost all OSKM-reprogrammable cells (**Fig. 3G-H**). Interestingly, the proliferation rate of the two systems was different as well, though both reprogramming combinations contained Myc. Based on the expression of proliferation gene signature, it was clear that OSKM-reprogrammable cells proliferate much faster than GETM-reprogrammable cells, while non-reprogrammable cells from both systems expressed these genes to a very low level (**Extended Data Fig. 3E**). This is in agreement with the proliferation rates of ESCs, whereby ESCs exhibit a very rapid doubling time of every 10-14 hours^46^.

In conclusion, these results illuminate the various fates, stage-specific markers and unique identifiers for each reprogramming system and strengthen the notion that each reprogramming process takes a distinct trajectory toward its own fate.

### Unique Methylation Dynamics for GETM and OSKM Reprogramming

One of the crucial aspects of nuclear reprogramming is the ability to erase the epigenetic landscape of the somatic nucleus and construct a new epigenetic profile that is similar to the target cells (e.g. ESCs for iPSCs and bdTSCs for iTSCs). One important epigenetic mark is DNA methylation, which allows chromatin condensation and silencing of specific loci along the genome^14^.

To assess the methylation landscape of somatic cells undergoing reprogramming to iTSCs and iPSCs, we applied the reduced representation bisulfite sequencing (RRBS) technique to capture the CpG methylation landscape of the cells as a representation for the global methylation changes. GETM and OSKM-reprogrammable cells from different time points (**Extended Data Fig. 1D**), as well as the parental fibroblasts, bdTSCs, iTSCs, ESCs and iPSCs were subjected to RRBS and analyzed.

We used the K-means algorithm to classify ∼130,000 genomic regions (blocks) shared amongst all samples during reprogramming to a TSC or pluripotent state, and generate 100 unique clusters where some of the clusters contained tiles that are specific to TSCs and ESCs, other to MEFs and the vast majority to reprogrammable cells (**Extended Data Fig. 4A**). Average DNA methylation levels per sample per cluster were then projected onto the first two principal components which generated gradual and time-dependent methylation dynamics for each reprogramming system with a clear ‘V’-shaped trajectory, where PC1 represents the OSKM trajectory and PC2 GETM trajectory (**Fig. 4A**). The accuracy of the time-dependent methylation trajectory in the two reprogramming systems was surprising, given that there is often poor correlation between methylation degree and gene activation, and the unique transcriptional profiles that characterize intermediate reprogrammable cells (**Fig. 2A and 2C**).

**Fig. 4.**
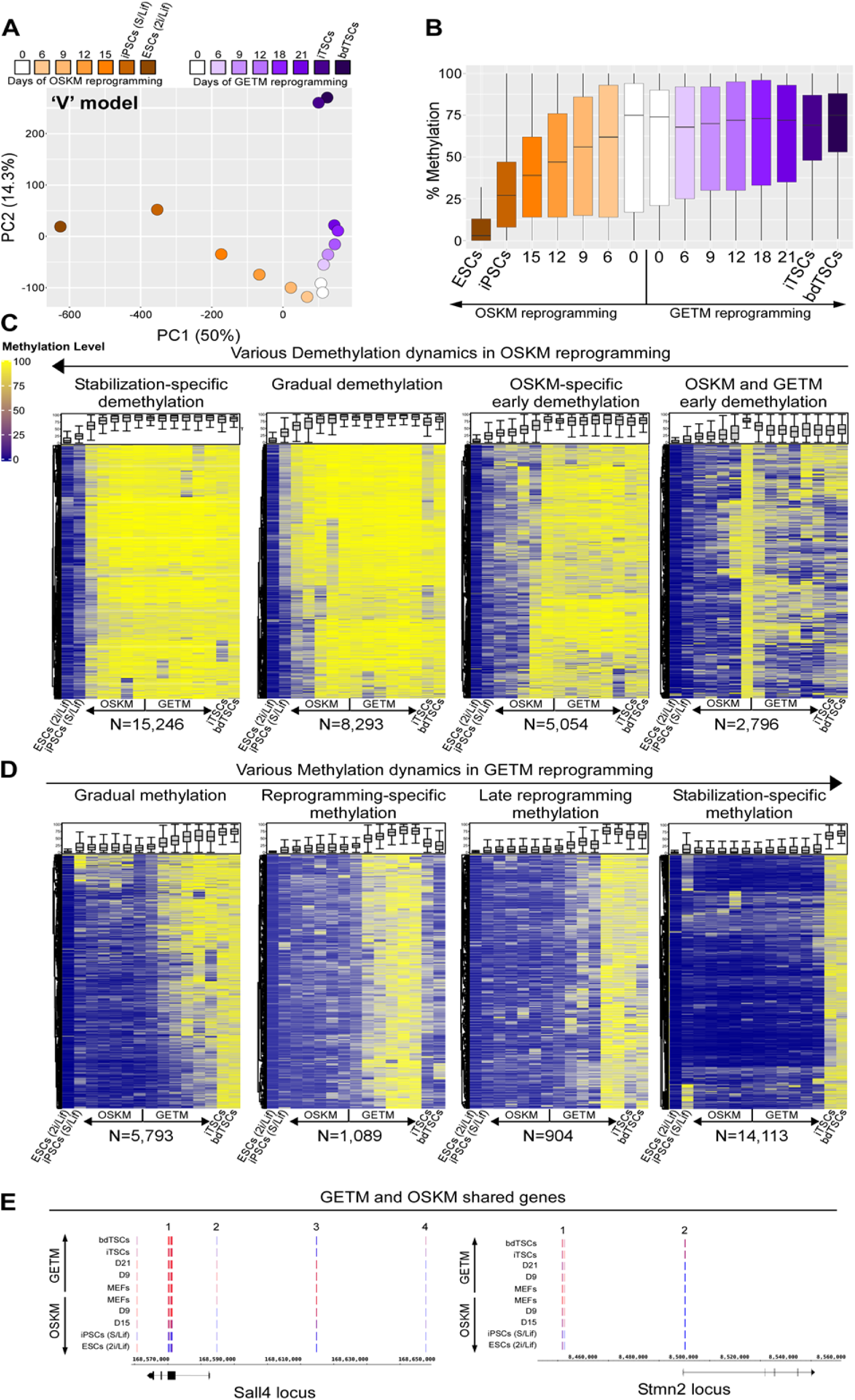
RRBS analysis demonstrates methylation specific dynamics between OSKM and GETM reprogramming. **(A)** Average bulk DNA methylation data of cells undergoing reprogramming toward pluripotency and TSC state projected onto the first two principal components. A clear V-like shape progression is observed separating GETM from OSKM reprogramming. **(B)** Boxplot of DNA methylation level across bulk samples during reprogramming towards both pluripotent and TSC states. OSKM reprogrammable cells exhibit an overall hypomethylation dynamics on CpG-enriched sites while GETM reprogrammable cells exhibit hypermethylation dynamics. **(C-D)** Heatmaps demonstrating the dynamics of DNA methylation alterations and patterns across bulk samples during reprogramming towards both pluripotent and TSC states, respectively. Each row represents one differentially methylated block for which there are at least one CpG with ≥10× coverage. While most of the patterns observed are specific to each reprogramming system, a few clusters show similar DNA demethylation initiated as early as day 6 in both systems. **(E)** DNA methylation level of genomic loci containing genes that are shared between pluripotent cells and TSCs.

One clear difference between OSKM and GETM reprogramming was the overall dynamic of methylation changes during the reprogramming process (**Fig. 4B**). While OSKM-reprogrammable cells predominately lose methylation on CpG-enriched sites during the reprogramming process, GETM-reprogrammable cells mostly gain methylation on CpG-enriched sites either in the middle or gradually until the end of the reprogramming process (**Fig 4B-D** and **Extended Data Fig. 4B-D**).

Mammalian placentas are unique in their methylation landscape as they contain regions in the genome that are highly methylated in gene bodies and regions that are only intermediately methylated (40=60%)^47^. In accordance with that, our unbiased analysis identified two unique clusters that contain intermediately methylated regions only in the final and stabilized iTSCs/TSCs (**Extended Data Fig. 4E**). These results suggest a unique mechanism of methylation/demethylation that occurs only when the core TSC circuitry is activated.

We then associated the various tiles from the different clusters to their neighboring genes and performed GO analysis using GREAT^48^ (**Supplementary Table 2**). Interestingly, while clusters associated with gradual loss of methylation in OSKM reprogramming involve genes that participate in the maintenance of the fibroblastic identity, apoptosis and multiple somatic cell properties (e.g. GO terms of actin cytoskeleton organization, focal adhesion, regulation of small GTPase mediated signal transduction, fat differentiation, muscle development, vasculogenesis and keratinocyte differentiation), the singular cluster among these which exhibits early demethylation in both systems is composed of genes that participate in somatic stem cell maintenance, immune system development and regulation of growth (**Supplementary Table 2**). While demethylation of regions that play a role in stemness and growth can be expected in the two systems, the identification of a set of genes that is enriched in the immune system, in methylation pattern and RNA in both systems, is intriguing (**Extended Data Fig. 3A-B and Supplementary Table 2**).

Clusters that are associated with gradual gain of methylation specifically in GETM reprogramming mostly include genes that negatively regulate metabolic processes involved in RNA production and transcription (e.g. RNA biosynthesis, macromolecule metabolic processes, nitrogen compound metabolic process). In accordance with their identity as extraembryonic cells, clusters that involve methylation only at the final step of the reprogramming process and in the fully reprogrammed cells are comprised of genes that are essential for embryo development, neuronal lineage development and somatic cell differentiation at large (**Supplementary Table 2**).

These data suggest that while GETM factors utilize DNA methylation to shut off all master genes important for executing the embryonic development program, OSKM first open these regions and subsequently regulate their expression by histone modifications (as will be discussed in the next section). Of special note is the neuronal lineage: While OSKM activates this program, GETM induces its silencing.

Given the fact that pluripotent cells and TSCs share many stemness genes (e.g. Sall4, Esrrb, Sox2, Lin28 etc. ^13, 42^), we next asked whether we can identify methylation differences in their regulatory elements during reprogramming and in the final cells. We selected 6 genomic loci that contain tiles for genes that are either specific to pluripotent cells (Slc15a1 and Tex19.2), specific to TSCs (Eomes and Bmp8b) or shared between the two cell types (Sall4 and Stmn2). Interestingly, only few tiles on regulatory elements (e.g. tile block number 2 and 3 in Sall4 locus) were methylated/hypomethylated similarly between iPSCs/ESCs and bdTSCs/iTSCs and different from MEFs, weakening the notion of widespread shared regulatory elements between the two cell types (**Fig. 4E**). Most tiles on regulatory elements were methylated oppositely between the two cell states (**Fig. 4E** **and Extended Data Fig. 4F-G**), indicating tight and cell type-specific regulation for each reprogramming process.

Taken together, these data suggest that although the acquisition of the final methylation landscape of GETM and OSKM reprogrammable cells is gradual and time-dependent, the methylation level and deposition is unique for each reprogramming process, even in genes that are expressed in both cell types.

### Chromatin Accessibility and Activity of Cells Undergoing Reprogramming to Pluripotent and TSC states Demonstrate Unique Chromatin Dynamics for each Reprogramming Process

One of the properties of reprogramming factors is their ability to open closed chromatin by recruiting chromatin remodelers and transcriptional machinery to heterochromatin^49, 50^. One approach to assess chromatin accessibility and activity is to employ Assay for Transposase-Accessible Chromatin using sequencing (ATAC-seq)^51^ in conjunction with chromatin immunoprecipitation and sequencing (ChIP-seq) for specific histone marks.

To assess which sites are open and active and which are closed during early stages of reprogramming to iPSCs and iTSCs, we collected cells (50,000 cells per replicate for ATAC-seq and 50 million cells per replicate for ChIP-seq for H3K4me2 and H3K27ac) at day 3, 6 and 9 of the reprogramming process to iTSCs and iPSCs (**Extended Data Fig. 1D**). As a control, we profiled chromatin accessibility and activity of the parental MEFs, the final reprogrammed cells (i.e. iTSCs and iPSCs) and their blastocyst-derived controls. We chose H3K27ac and H3K4me2 because H3K27ac marks both active promoters and distal enhancers while H3K4me2 is enriched in *cis*-regulatory regions, particularly in promoters, of transcriptionally active genes but more importantly it also marks genes primed for future expression^52, 53^. Overall, we analyzed 170,658 peaks for ATAC-seq, 498,376 for H3K27ac and 770,274 for H3K4me2. These experiments allowed us to map the regions that are open and active early in the reprogramming process and those that are primed to be active later as well as closed regions in each reprogramming system.

In accordance with the transcriptome and methylome results, PCA on datasets of chromatin accessibility (ATAC-seq) and activity (i.e. H3K27ac and H3K4me2) revealed two separate ‘V’-shaped trajectories distinguishing OSKM from GETM reprogramming, already at the beginning of the process (**Fig. 5A-C**). These results suggest that OSKM and GETM remodel the chromatin at different regions. Following this observation, we asked whether the distribution of the various peaks along the genome (i.e. promoters, exons, introns, UTRs, TSS and intergenic) is unique to each reprogramming system or a global reprogramming phenomenon. We found that while the location of the various peaks along the genome is mostly different between the two reprogramming systems (**Extended Data Fig. 5A-C**), the distribution of the peaks is very similar (**Fig. 5D-F**).

**Fig. 5.**
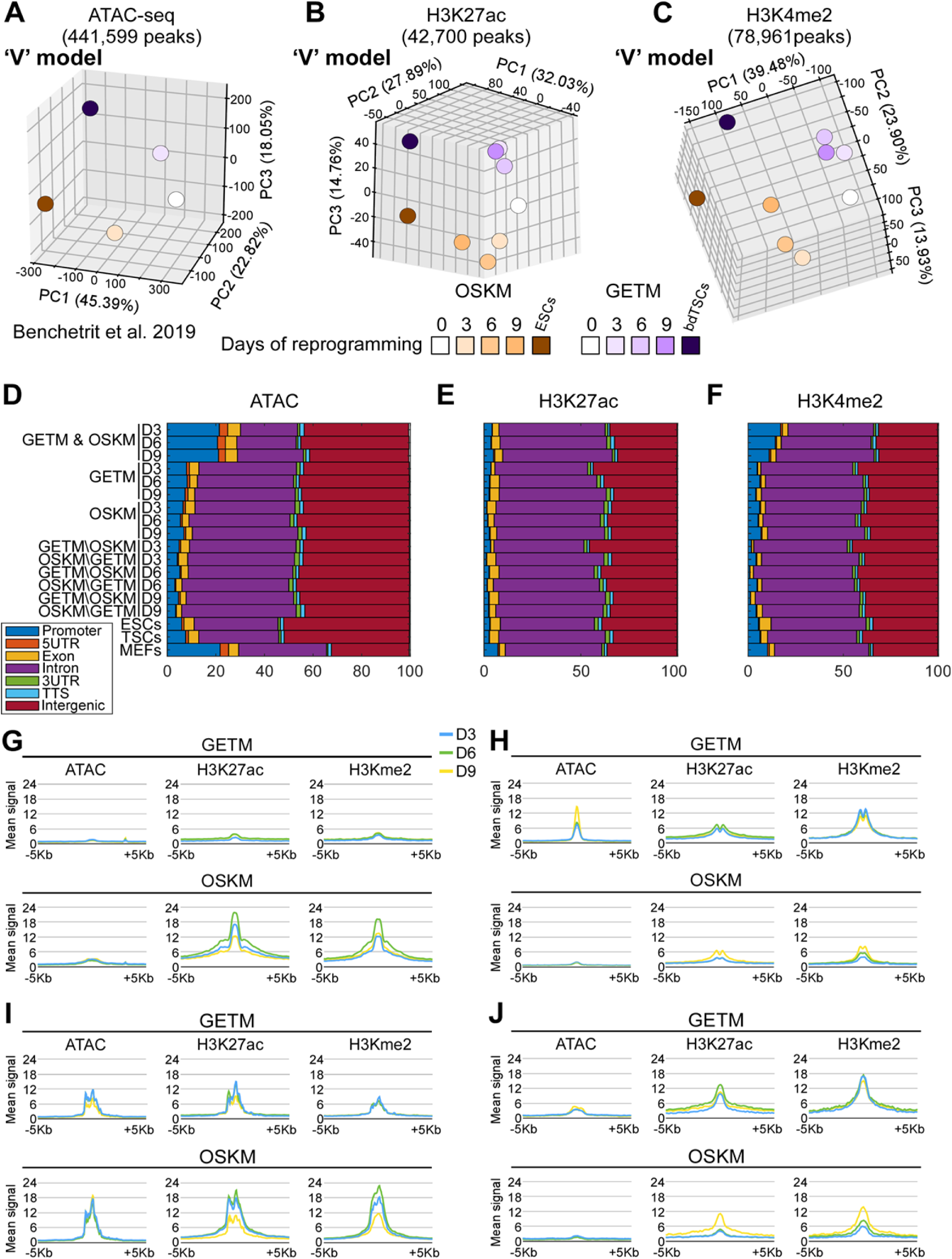
Chromatic accessibility and activity during GETM and OSKM reprogramming demonstrating a ‘V’-shaped behavior. **(A-C)** Top 3 PCA components of the Z-scores of ATAC-seq (A), H3K27ac (B) and H3K4me2 peaks (C). Peaks were clipped to range [0, 500] and filtered by length (>=500bp). Replicates were merged by taking the mean peak height. **(D-F)** Genomic annotations of ATAC-seq peaks (D), H3K27ac peaks (E) and H3K4me2 peaks (F). Shown are the fraction of various genomic annotations (Promoter, Exons, Introns, etc) among peaks. Genomic regions accessible in both GETM and OSKM conditions (D, top three rows) are enriched for promoter regions, compared to GETM or OSKM regions (below). GETM and OSKM mark regions accessible in those conditions, excluding MEF peaks. Below are cell-type specific accessible regions such as “GETM\OSKM D3”, which includes GETM Day 3 peaks not accessible in OSKM Day 3. In addition to Promoter regions (blue), most accessible regions fall within Intronic regions (purple) and Intergenic regions (red). **(G-J)** Mean ATAC-seq, H3K27ac and H3K4me2 at ±5Kb surrounding OSKM or GETM H3K27ac peak locations, for Day 3 (blue), Day 6 (green), and Day 9 (yellow). (G) OSKM-specific H3K27ac signal is strongest at Day 6, and is accompanied by matching H3K4me2 signal, but with no dynamic change in DNA accessibility. (H) Same for GETM ATAC-seq peaks. These regions are already marked by H3K4me2 in Day 3, and gain accessibility over time. These genome regions also show H3K27ac and H3K4me2 enrichments for later OSKM stages. (I) Same for OSKM ATAC-seq peaks. (J) Same of GETM H3K27ac peaks. These peaks show gradual increase in ChIP-seq signal even following OSKM induction (below).

Interestingly, while shared OSKM and GETM ATAC-seq peaks and, to a lesser extent H3K4me2 peaks, exhibit a significant enrichment in promoters and exons, peaks that are unique to each reprogramming process are mostly localized to introns and intergenic regions (**Figs. 5D and 5F**). EnrichR GO analysis on the genes associated with the shared open promoters suggested that many of these active promoters are associated with response to the lentiviral infection itself (p≤0.001). No significant differences in the distribution of H3K27ac is found between the various samples. These data suggest that the reprogramming process at large follows the same rules of genomic remodeling (i.e. beginning with robust opening of intronic and intergenic regions in conjunction with promoter closing), but each reprogramming system remodels the chromatin at different loci along the genome in accordance with its final cellular fate.

We then classified the various peaks (mean ATAC-seq, H3K27ac and H3K4me2 at ±5Kb) based on their behavior in the two reprogramming processes (**Extended Data Fig. 6A-D**). We identified 4 distinct patterns: (1) 1,605 genomic regions which appear predominantly in OSKM reprogramming in which their H3K27ac signal is typically the strongest at day 6 and is accompanied by a matching H3K4me2 signal, but with no dynamic change in DNA accessibility (**Fig. 5G**). (2) 1,716 GETM-specific regions that are marked by H3K4me2 and H3k27ac already at day 3 but chromatin accessibility is gained only later in the process (i.e. day 9). Intriguingly, a mirror image can be seen in these regions during OSKM reprogramming. There, chromatin accessibility is mildly open and remained unchanged but a significant increase in H3K27ac and H3K4me2 signals is observed at later stages of reprogramming (**Fig. 5H**). (3) 2,848 regions that are open and active in both reprogramming systems but lose activity (i.e. H3K27ac and H3K4me2 signal) over time exclusively in OSKM reprogramming (**Fig. 5I**). (4) 464 genomic regions that are open in both reprogramming processes, but while in GETM reprogramming they are active all along the process, in OSKM reprogramming they gain activity exclusively at later stages (**Fig. 5J**). We used GREAT to test these genomic regions for enriched annotations. We found that group 1 is associated with cellular response to leukemia inhibitory factor (p≤8.12e-23), Ras guanyl-nucleotide exchange factor activity (p≤5.0e-9) and represents a phenotype of embryonic lethality between implantation and placentation in the mouse (p≤1.5e-12). Group number 2 is associated with cell migration and motility (p≤2.6e-20), cell adhesion (p≤4e-14), extracellular matrix (p≤1.0e-8), insulin-like growth factor binding (p≤2.7e-8) and heparin binding (p≤3.0e-7), all relevant to trophoblast differentiation and placentation. Interestingly, once again, this group of genes is significantly enriched in cells of the immune system, giving rise to the GO term of mouse phenotype of autoimmune response (p≤3.4e-13). These results suggest a mechanism by which GETM induce a TSC fate by gradually opening and activating trophoblast-specific regions that are important for TSC function. In contrast, OSKM do not change the accessibility of these regions, which remain mildly open, but then gradually activate them, which explains the small fraction of differentiated trophoblast cells present in OSKM reprogramming.

**Fig. 6.**
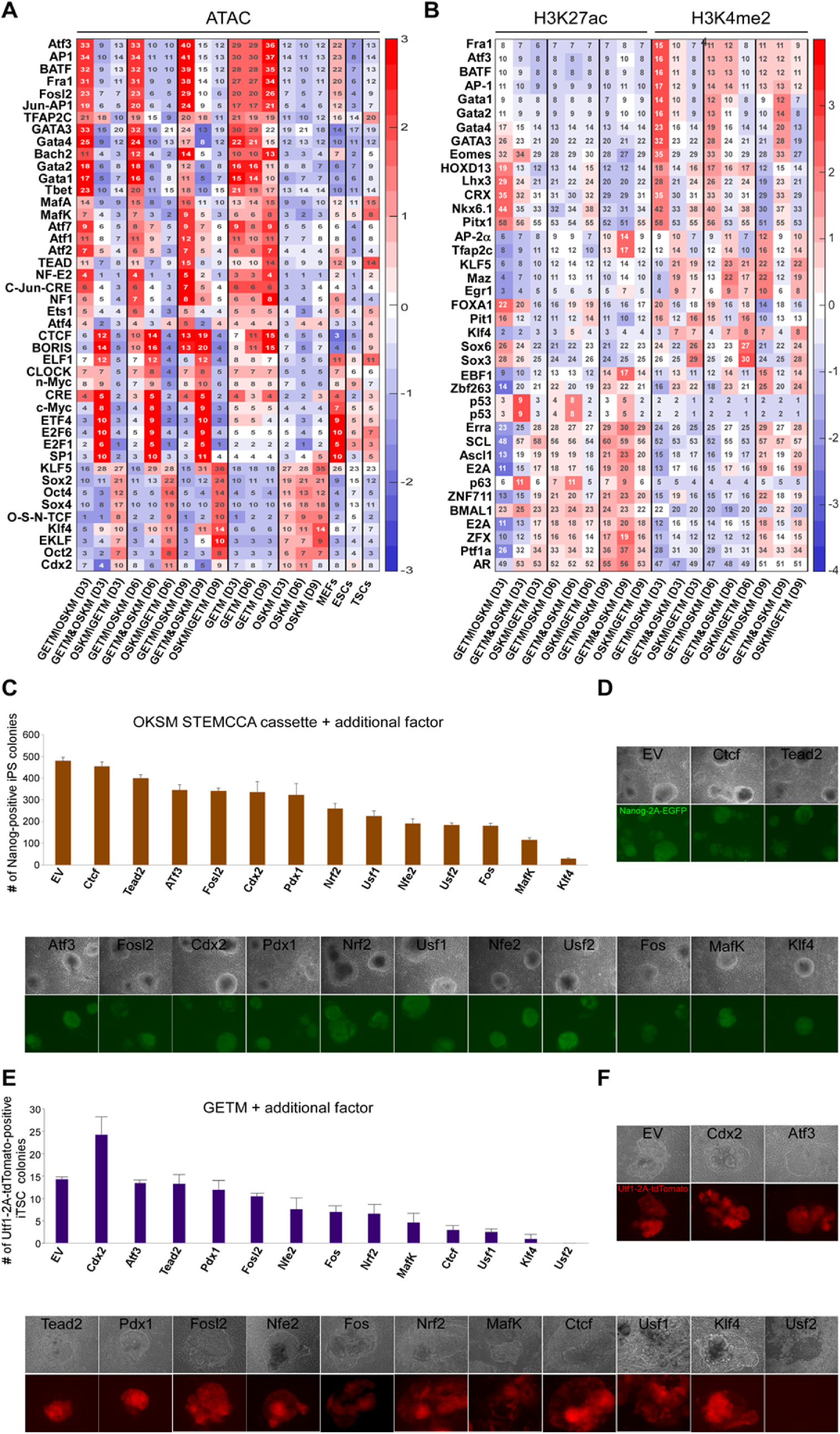
Motif enrichment and the effect of their corresponding transcription factor on OSKM and GETM reprogramming. **(A)** Heatmap showing motif enrichment among ATAC-seq peaks. For each row (motif) and each column (condition-specific ATAC-seq peaks) we calculated the percent of peaks containing it (shown numbers). Subsets of peaks include GETM-only peaks (GETM\OSKM), joined peaks (GETM&OSKM), and OSKM-only peaks (OSKM\GETM) for each time point (Days 3, 6, 9). Also shown are joined sets of GETM and OSKM peaks for each day, as well as MEF, ESC and TSC peaks. Each motif/condition is color-coded based on relative motif enrichment (Z-scores) compared to all conditions. Only motifs with enrichment greater than 2.5 standard deviations (Z>2.5) are shown. **(B)** Heatmap showing motif enrichment among H3K27ac and H3K4me2 peaks. For each row and each column, we calculated the percent of peaks containing it (shown numbers). Subsets of peaks include GETM-only peaks (GETM\OSKM), joined peaks (GETM&OSKM), and OSKM-only peaks (OSKM\GETM) for each time point (Days 3, 6, 9). **(C-D)** BYKE MEFs were infected with dox-inducible OKSM STEMCCA cassette plus additional factor as depicted. The cells were reprogrammed for 8 days and then weaned of dox for additional 5 days. Nanog-2A-EGFP-positive colonies were counted (C) and imaged (D). EV refers to empty vector control. **(E-F)** BYKE MEFs were infected with dox-inducible GETM factors plus additional factor as depicted. The cells were reprogrammed for 21 days and then weaned of dox for additional 10 days. Utf1-2A-tdTomato-positive colonies were counted (E) and imaged (F). EV refers to empty vector control.

Group number 3 contains regions that are related to platelet-derived growth factor receptor signaling pathway (p≤1.8e-5), ERBB signaling pathway (p≤8.5e-5), epidermal growth factor receptor signaling pathway (p≤1.5e-4), with a mouse phenotype of placental labyrinth hypoplasia (p≤2.6e-5). Group number 4 is associated with genes involve in cell motility and cell migration (p≤8.1e-8), focal adhesion (p≤1.3e-14) and actin cytoskeleton (p≤3.6e-13), and abnormal extraembryonic tissue morphology (p≤3.1e-10).

Overall, OSKM and GETM factors open and activate regions that are essential for their function (e.g. Wnt signaling for OSKM and cell migration, motility and reaction to heparin and insulin for GETM) as well as regions that are important for the induction of MET and cellular transformation (e.g. focal adhesion and actin cytoskeleton).

Next, we subtracted all the peaks that were overlapped with MEFs to identify transcription factor binding sites that are enriched in peaks (ATAC-seq, H3K27ac and H3K4me2) associated with each reprogramming process at various reprogramming time points. (**Fig. 6A-B** **and Supplementary Figs. 5A-C and 6E-F**). We observed a highly significant P-value for binding motifs of OSK factors in OSKM reprogramming peaks and GET binding motifs in GETM reprogramming peaks, supporting our analysis (**Figs. 6A-B, Supplementary Figs. 5A-C and 6E-F**). Interestingly, the binding sites of the AP1/CREB/ATF families of proteins, which act as somatic cell identity safeguards and block the reprogramming process to pluripotency^41^, are significantly more enriched in GETM reprogramming peaks compared to OSKM reprogramming peaks (**Figs. 6A-B and Extended Data Fig. 5A-C**). In contrast, regions that are open in the fibroblasts and closed upon reprogramming, the binding sites of the AP1/CREB/ATF family of proteins are significantly more enriched in OSKM reprogramming compared to GETM reprogramming (**Extended Data Fig. 6G-H**). These results explain the potent ability of OSKM to erase somatic cell identity, as well as the continued presence of MEF-like cells in GETM reprogramming even at day 12 of the process (**Fig. 3A-B** and **Extended Data Fig. 3A**). Besides the expected Gata, Tfap2c and Eomes/Tbet motifs and the binding sites for AP1/CREB/ATF families, GETM-specific peaks were enriched with genes involved in oxidative stress response such as Nrf2^54^, Nfe2^54^, MafK^54^ and Bach1/2^55^ (**Fig. 6A-B**, **Supplementary Figs. 5A-C and 6E)**. In contrast, OSKM-specific peaks were enriched with pluripotency binding sites such as Klf, Sox, Oct and Nanog as expected, but also with genes involved in neuronal differentiation, such as E2A^56^, Ascl1^57^, and with trophoblast such as Cdx2 and Znf263 (**Figs. 6A-B and Supplementary Figs. 5A-C and 6F)**, explaining the generation of trophoblast-like cells and neuronal fate observable in OSKM reprogramming (**Figs. 2H and 3B**). In agreement with their role as reprogramming factors, GETM and OSKM shared peaks were enriched with genes that are important for remodeling the chromatin such as Ctcf^58^, BORIS^59^, E2f6^60^, Elf1^61^, Usf1/2^62^ and YY1^63^.

We then chose 13 factors whose binding sites were significantly enriched in either GETM reprogramming (Nrf2, Nfe2, Fos, MafK, Atf3, Fosl2, Tead2) or OSKM reprogramming (Klf4, Cdx2, Pdx1) or shared between the two systems (Ctcf, Usf1, Usf2,). We performed reprogramming experiments to iPSCs and iTSCs with BYKE cells transduced with either OSKM or GETM, together with an empty vector (EV) control or with one of the 13 selected factors (**Fig 6C-F**). Strikingly, all examined factors (besides Cdx2 in GETM) either hindered the reprogramming process or had a mild effect in both systems, suggesting that both OSKM and GETM initially open regions that are highly regulated by somatic identity safeguards that counteract the reprogramming process at large (**Fig. 6C-F**).

Surprisingly, a very strong reprogramming inhibition was noted when Klf4 was overexpressed in both GETM and OSKM reprogramming. As Klf4 is relatively highly expressed in MEFs, it is tempting to speculate that overexpression of Klf4 on top of OSKM alters the stoichiometry of the reprogramming factors and counteracts reprogramming by maintaining fibroblastic identity (**Fig. 6C-F**). Another very strong reprogramming blocker that we found, especially for iTSC reprogramming, is Usf2, which is a known tumor suppressor and Myc inhibitor^64^. Moreover, Usf2 is a strong regulator of iron metabolism and oxidative stress response^65, 66^, proposing an explanation as to why Usf2 has a stronger effect on iTSC reprogramming.

In contrast to the global reprogramming blockers mentioned above, Ctcf, which significantly hindered the reprogramming to iTSCs, only mildly affected the reprogramming to iPSCs (**Figs. 6C and 6E**). As Ctcf is highly expressed in iTSCs/TSCs and acts as a very important chromatin insulator that controls gene expression^67^, this result emphasizes the importance of retaining normal levels of Ctcf for the induction of the TSC fate. As expected, Cdx2 facilitated the reprogramming to the TSC state and hindered reprogramming to the pluripotent state.

We then analyzed the binding sites of closed regions; peaks that were open in MEFs and disappeared during the reprogramming process with GETM or OSKM (**Extended Data Fig. 6G-H**). Interestingly, while OSKM closed peaks are enriched with binding sites of AP1/CRE/ATF family, TEAD, Pdx1, RUNX and Mef2a, indicating the initial loss of the fibroblast identity, GETM closed peaks are enriched as well with RUNX and TEAD but also with interferon response genes such as STAT5, ISRE, RXR and IRF1/2 and apoptosis-related genes such as p53 and p63. This might suggest that GETM overcome viral infection-induced apoptosis by closing regions that control interferon response genes and master regulators of cell death.

Next, we sought to determine whether the regions that begin to open up via GETM and OSKM during the initial phase of reprogramming are active or not. Using scatter plots, we probed all ATAC-seq peaks from both GETM and OSKM reprogramming (**Extended Data Fig. 7A**). In accordance with the PCAs and Venn diagrams, the vast majority of the peaks were unique to each reprogramming process. We then plotted all the H3K4me2 peaks on top of the ATAC-seq peaks (**Extended Data Fig. 7B**) and performed GO analysis on OSKM or GETM-specific peaks using EnrichR (**Extended Data Fig. 7C-D**). By comparing OSKM and GETM H3K4me2 peaks, we were able to focus on all the unique regions that are remodeled by OSKM and by GETM, as any global regions that are involved in the identity of the fibroblast or are important for reprogramming at large will overlap between the two reprogramming systems. Remarkably, besides regions that are involved in the regulation of epithelial cell migration, analyzing OSKM-specific peaks revealed significant enrichment for regions important for the development of the heart (e.g. GO terms of regulation of heart contraction and regulation of cardiac conduction), the first and arguably most crucial organ to form during embryogenesis (**Extended Data Fig. 7C**). Moreover, a significant enrichment was found for the formation of the brain and liver as well (e.g. GO terms of neuron projection maintenance, axon guidance and determination of liver asymmetry, **Extended Data Fig. 7C**). In contrast, GETM-specific H3K4me2 peaks are enriched for regions that involve metabolic processes and proliferation, as well as regions that are essential for trophoblast function such as migration and attraction of blood vessels (e.g. GO terms of regulation of blood vessel cell migration, regulation of cell migration in angiogenesis, **Extended Data Fig. 7D**).

**Fig. 7.**
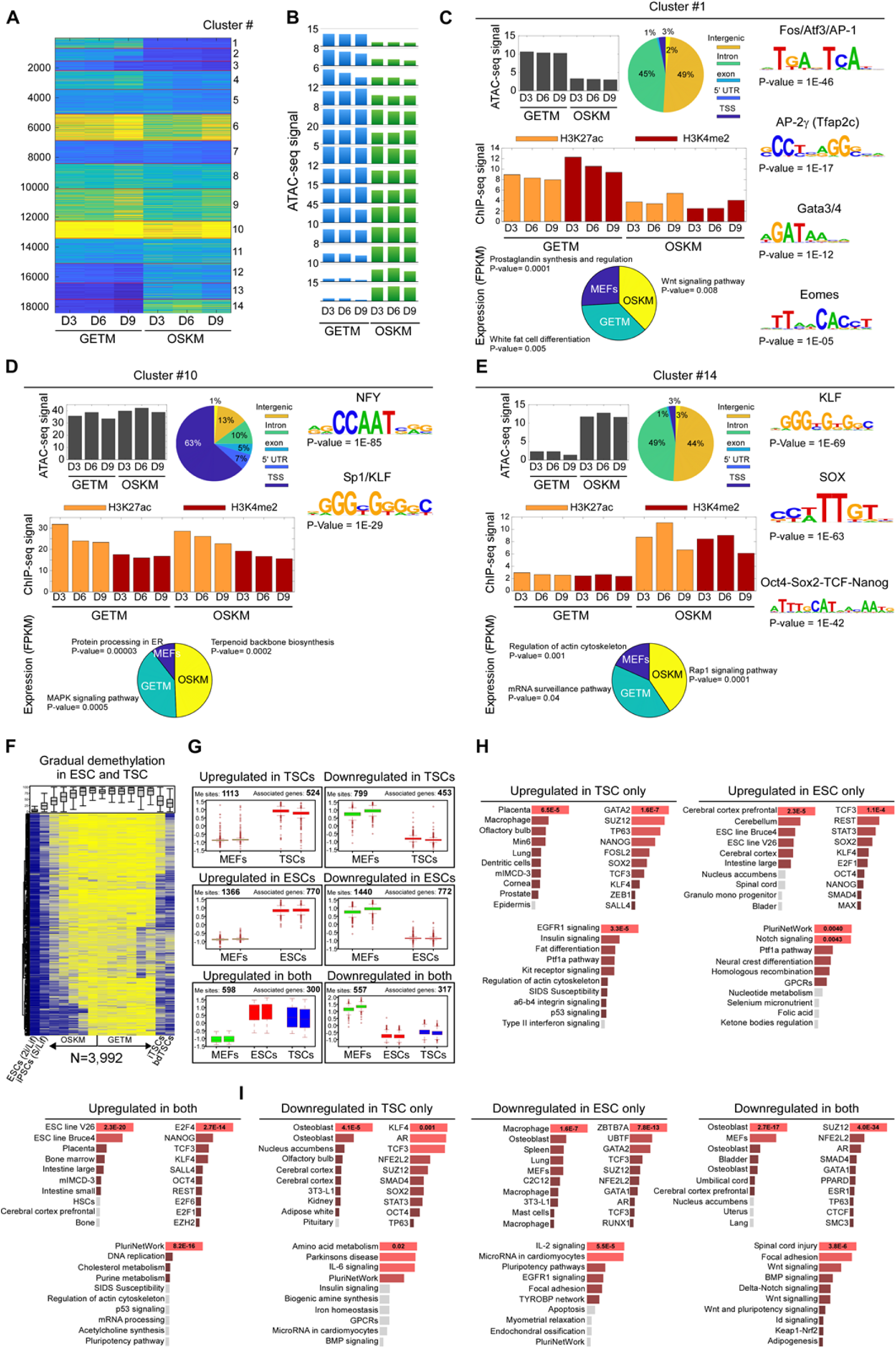
Data integration of DNA methylation, gene expression and chromatin accessibility and activity during the reprogramming process toward pluripotent and TSC states. **(A-B)** Clustering of 18,420 GETM and OSKM ATAC-seq peaks from days 3, 6, 9 into 14 clusters is shown as a heatmap (A) or barplot of mean ATAC-seq signal per cluster (B). **(C)** Cluster #1 is mostly composed of distal (Intergenic and Intronic) GETM-specific peaks, enriched for AP, GATA and Eomes motifs, and near GETM-expressed genes. Shown are mean ATAC-seq signals (top left), analysis of their genomic annotations (pie chart, center), enriched transcription factor motifs (right panel), average ChIP-seq signals of H3K27ac and H3K4me2 following GETM and OSKM induction (middle panel), and a pie chart for RNA expression levels and GO term for genes that are associated with each cluster ATAC-seq peaks and exhibit the highest expression levels in MEFs (blue), or GETM (green) or OSKM (yellow, Bottom panel). **(D)** Same for cluster 10, enriched for highly accessible promoter peaks. **(E)** Same for cluster 14, with regions that are highly accessible following OSKM, enriched for distal regions with KLF, SOX and Oct4 motifs, and are associated with OSKM expressed genes. **(F)** A heatmap of differentially methylated blocks with DNA demethylation during the final states of reprogramming to both pluripotent and TSC states. Each row represents one out of 3992 blocks of DMBs. **(G)** Boxplots of relative expression of differentially expressed genes that are associated with each individual block of DNA methylation. A significant negative correlation between DNA methylation and expression is observed in both reprogramming systems in which DMBs are associated with genes upregulated in both systems such as Esrrb, Sox2, Sall4, Dppa4 and Zfp42. A significant positive correlation is also observed in both reprogramming systems in which DMBs are associated with MEFs genes downregulated in both systems such as Acta2, Col5a1, Col5a2, Runx1 and Runx2. **(H-I)** Enrichr Mouse gene atlas and KEGG pathways analysis of significantly over-represented genes that are either upregulated (H) or downregulated (I) for each depicted group in (G).

We then plotted the active histone mark H3K27ac on top of the ATAC-seq peaks (**Extended Data Fig. 7E**). Interestingly, we noted that while OSKM-specific ATAC-seq peaks tend to lose H3K27ac during the reprogramming process, GETM-specific peaks gain H3K27ac (**Extended Data Fig. 7F**). OSKM-specific H3K27ac peaks are mainly enriched within regions that play a role in neuron development and Wnt and calcium signaling pathways (**Extended Data Fig. 7G**), while GETM-specific peaks are enriched within regions that involve the regulation of MAPK activity, response to reactive oxygen species (ROS) as well as various metabolic processes (e.g. GO terms of metal ion homeostasis, Camp-mediated signaling and protein phosphorylation, **Extended Data Fig. 7H**).

Overall, these results agree with the transcriptome and methylation data, suggesting that OSKM initially open (ATAC-seq peaks) and define (H3K4me2 peaks) regions along the genome that are important for their differentiation potential but then eliminate their activity by removing the active mark, H3K27ac. In contrast, GETM specifically open and activate regions that are essential for trophoblast function as the process progresses, while closing and methylating regions which participate in the embryonic development program.

### Data Integration Analysis Illuminates Key and Unique Aspects for the Induction of Pluripotency and TSC State

We then performed data integration analysis to correlate gene expression to chromatin accessibility and activity (**Figs. 7A-E and Extended Data Fig. 8A-K**) as well as to methylation (**Figs. 7F-I**) To that end, we initially performed cluster analysis on 18,420 GETM and OSKM-specific ATAC-seq peaks. This gave rise to 14 clusters (**Fig. 7A-B**) that show unique patterns of chromatin accessibility, activity, peak distribution and DNA binding motifs in the two reprogramming systems. Then, we associated the ATAC-seq peaks of each cluster to their neighboring genes and tested whether their expression is highest in GETM reprogrammable cells, OSKM reprogrammable cells or in MEFs (depicted as a pie graph in the bottom part of each cluster). Finally, for each group of genes (i.e. highest in GETM, OSKM or MEFs) we performed GO annotation.

Clusters 1-4 are GETM-specific clusters, as they harbor genomic regions with a higher chromatin accessibility and activity in GETM samples compared to OSKM samples (**Figs. 7C and Extended Data Fig. 8A-C)**. While clusters 1,2,3 are enriched for binding motifs of the GETM reprogramming factors Gata3, Tfap2c and Eomes as well as Tbx6 and FOS/Atf3/AP-1, cluster 4 is enriched for binding motifs for AP-1 and for the master TSC regulator, TEAD ^68^. Moreover, while clusters 1,2,3 contain mostly intronic and intergenic ATAC-seq peaks, cluster 4 encompasses a large fraction of transcription start site (TSS) ATAC-seq peaks. GO annotation analysis revealed that clusters 1,2,3 include genes that are associated with fat differentiation (p≤0.005), p38 MAPK pathway (p≤0.001) and Glycogen metabolism (p≤0.0002), while cluster 4 contains genes involved in apoptosis (p≤0.0007).

In accordance with that, it has been shown that lipid droplets formation and glycogen storage are crucial for proper trophoblast function and that p38 MAPK pathway controls the invasiveness capability of trophoblastic cells^69–71^. Genes that are upregulated in clusters 1,2,3 to the highest level in OSKM reprogrammable cells are related to Wnt signaling (p≤0.008), IGF-1 pathway (p≤0.01) and cardiocyte differentiation (p≤0.0004), while cluster 4 is enriched with genes involve in BMP signaling pathway (p≤0.0001). Interestingly, regions that control Wnt and IGF-1 pathways which are implicated in pluripotency maintenance ^72^ are also more activated in GETM reprogramming but the associated genes are higher in OSKM reprogramming suggesting that these regions are negatively regulated by GETM. In contrast, genes that are expressed to the highest level in the parental MEFs in clusters 1,2,3 are connected to prostaglandin synthesis (p≤0.0001), integrin binding (p≤0.00001) and Tgf-β signaling (p≤0.004), while cluster 4 is enriched with genes that involve in focal adhesion (p≤7e-12), all implicated in fibroblastic identity maintenance^73, 74^ and are negatively regulated by GETM factors.

Clusters 5-11 are GETM and OSKM shared clusters as they harbor genomic loci with either high or low chromatin accessibility and activity in both reprogrammable cells (**Figs. 7D and Extended Data Fig. 8D-I**). Clusters 6,8,9,10, are highly enriched with strong peaks around the TSS and as such, contain binding sites for transcription factors that are implicated in transcription initiation, such as NFY, SP1/KLF, ETS, E2A and Oct2^75–77^. Intriguingly, genes that are upregulated in these clusters to the highest level in GETM reprogramming are implicated in mRNA processing (e.g. mRNA catabolic process (Pv ≤ 5.0e-9), mRNA processing (p≤5.0e-8), spliceosome (p≤9.1.0e-10)), while genes that exhibit the highest levels in OSKM reprogramming involve mitotic cell cycle regulation (p≤4.4e-11), estrogen signaling (p≤0.00002) and insulin signaling (p≤0.0001). These processes are general and characterize both states, however, OSKM reprogramming also upregulated genes implicated in terpenoid backbone biosynthesis (p≤0.0002) which is involved in the conversion of the pluripotency primed state to naïve state^78^. Genes with the highest expression levels in MEFs participate in protein processing in ER (p≤0.00003), EGFR1 signaling pathway (p≤0.000007), cyclic nucleotide catabolic process (p≤0.00007) and cell junction assembly (p≤0.000002). In contrast, clusters 5,7,11, which are also shared between OSKM and GETM reprogramming but enriched with smaller peaks that are located around the TSS as well as in intergenic and intronic regions, contain binding sites for transcription factors that are key drivers of the fibroblastic identity (Zeb1, Tcf12, Tbx5 and Sox6) and more significantly, with binding sites for the insulator gene Ctcf. In agreement with that, GO analysis for genes that are highest expressed in MEFs in these clusters exhibited gene ontologies of extracellular matrix organization (p≤1.6e-10) and integrin binding (p≤0.00001). In contrast, genes that are expressed to the highest levels in GETM reprogrammable cells are involved in RNA processing (p≤0.0001), translation (p≤7.4e-9) and regulation of cytokines biosynthesis (p≤0.0001), while genes that are highest in OSKM reprogrammable cells are enriched for axonal transport (p≤0.0002), negative regulation on bone remodeling (p≤0.00007) and ubiquitin conjugation binding (p≤0.000008), suggesting the initial opening of the different lineages.

Clusters 12,13,14 are OSKM-specific with open and active peaks in OSKM reprogrammable cells that are enriched for OSK reprogramming factor binding sites, Oct, Sox, Klf as well as for Nanog (**Figs. 7E and Extended Data Fig. 8J-K**). Similar to GETM-specific clusters, these OSKM-specific clusters are also enriched with peaks that are mostly localized to intronic and intergenic regions. Genes that are associated with these peaks and harbor the highest expression level in OSKM reprogrammable cells are involved in Rap1 signaling pathways (p≤0.0001), positive regulation of the non-canonical Wnt signaling pathway (p≤0.00009) and neuron development (p≤0.00004), all implicated in neuronal cell activity^79, 80^, further explaining why OSKM reprogramming can induce neuronal fate. Genes that are expressed to the highest levels in GETM reprogrammable cells play a role in ERBB signaling pathway (p≤0.000002), glycogen metabolism (p≤0.006) and mRNA surveillance pathway (p≤0.04), processes which are important for trophoblast differentiation. Finally, genes that are associated with these peaks but exhibit the highest levels in MEFs are implicated in regulation of actin cytoskeleton (p≤0.001) and regulation of cell migration (p≤3.4e-10), all important for normal fibroblastic function.

In conclusion, these data describe how GETM reprogramming differs from OSKM reprogramming in the induction and dynamics of various signaling pathways, metabolomic processes and well as in their ability to erase the somatic identity and to induce alternative cell fates during reprogramming. Moreover, it demonstrates that from the onset of the reprogramming process, GETM reprogramming is directed toward the TSC fate and activates various processes and pathways that are essential for TSC maintenance and differentiation.

We then sought to correlate gene expression to methylation. We focused on one interesting cluster in which demethylation occurs in most regions of both systems only at the final step of the reprogramming process (**Fig. 7F**). We associated the 3,992 tiles of this cluster to their neighboring genes and examined their expression in ESCs and TSCs (**Fig. 7G**). We identified 525 genes that are upregulated and 453 genes that are downregulated specifically in TSCs when compared to MEFs. 770 genes are upregulated and 772 genes are downregulated specifically in ESCs, while 300 genes are upregulated and 317 genes are downregulated in both ESCs and TSCs when compared to MEFs.

GO annotation analysis revealed that genes that are upregulated specifically in TSCs play a role in EGFR1 signaling, insulin signaling and fat differentiation and contain transcription factor binding sites of GATA2, SUZ12 and TP63, all important players in trophoblast formation and differentiation. Indeed, Jensen tissue analysis associated these genes with the placenta with the most significant P-value (p≤6.5e-5, **Fig 7H**).

Analysis of the genes that are specifically upregulated in ESCs identified pluripotency and notch signaling pathways as the most significant biological processes, with transcription factor binding sites that are enriched for many pluripotency genes such as Tcf3, Rest, Stat3, Sox2, Klf4, Nanog and Oct4. Interestingly, Jensen tissue analysis associated these genes with Cerebral cortex prefrontal, cerebellum as well as ESC lines (**Fig. 7H**). Genes that are upregulated in both ESCs and TSCs are involved in stemness at large, as pluripotency network and DNA replication were identified as the most significant biological processes and ESC lines and placenta the most significant tissues. In accordance with that, E2F4 and to a lesser extent other E2F family members, which play a key role in stem cell proliferation ^81^, are found to be the most significantly enriched binding sites in these tiles, in addition to binding motifs of pluripotency genes such as Nanog, Tcf3, Klf4, Sall4 and Oct4 (**Fig. 7H**). When we analyzed the genes that are downregulated either in TSCs, ESCs or both, they were all implicated in fibroblastic identity and function, ranking osteoblasts, MEFs and macrophages as the most significant cell types (**Fig. 7I**). In accordance with the downregulation of these genes, their associated tiles were enriched with binding motifs of factors known to induce strong transcriptional repression such Klf4, AR, Zbtb7a, UBTF, Suz12 and Nfe2l2, suggesting how these hypomethylated regions are associated with gene repression.

Taken together, this analysis allowed us to examine the final stage of demethylation that occurs in both GETM and OSKM reprogramming, which characterizes the stabilization stage of both cell types. Surprisingly, this final stabilization step still involves the erasure of the fibroblastic identity.

### Genomic Stability Analysis during OSKM and GETM Reprogramming Reveals Equivalent Frequency of Copy Number Variations (CNVs) during the Initial Phase of Reprogramming

One of the characteristics of TSCs is a unique methylation landscape that allows the activation of many repetitive elements within the trophoblast genome^82^. This unique property is believed to induce genomic instability^83, 84^ and indeed, multiple genomic aberrations are found in both iTSCs and bdTSCs following prolonged culture^13^. Therefore, we next aimed to understand whether GETM activation induces genomic instability already at the onset of the reprogramming process toward the TSC state. Since Myc is a known driver of genomic instability^85^, we reprogrammed fibroblasts into iTSCs by GET or GETM and to iPSCs by OSK or OSKM as a control. Cells were collected for copy number variation (CNV) analysis immediately following the infection (day 0), at day 3 of reprogramming and at day 6 of reprogramming. In addition, we examined ten iPSC clones and two previously characterized partially reprogrammed cells^23^ as reference cells. All genomic reads were aligned against the parental MEF genome. As can be seen in Figure S7L, while many CNVs were identified in one of the two partially reprogrammed iPSC clones and few CNVs in chromosome 1 or 8 in three out of ten fully reprogrammed iPSC clones, this analysis could not identify a significant amount of CNVs in the initial phase of both OSK/M and GET/M reprogramming (**Extended Data Fig. 8L**).

As the sensitivity of bulk whole-genome sequencing is limited and does not allow the detection of CNVs at single cell resolution, further examination is needed to fully address this question. However, we can confidently conclude that substantial genomic instability is not induced by GETM at the initial phase of reprogramming.

## Discussion

Nuclear reprograming by defined factors is a powerful tool in understanding cellular plasticity and cell fate decision^86^ and for the generation of various cell types from somatic cells. Since the discovery of iPSCs by Takahashi and Yamanaka in 2006^17^, the reprogramming process of fibroblasts to iPSCs by the OSKM factors has been investigated extensively by many groups^22–37^. In contrast, the reprogramming process of fibroblasts to iTSCs by GETM, described for the first time in 2015^13, 16^, has never been done before.

Considering the fact that pluripotency and trophectoderm fates arise simultaneously during blastocyst development, we hypothesized that performing a parallel and comparative multi-omics analysis on both reprogramming processes concomitantly will yield knowledge that cannot be revealed otherwise, such as when each reprogramming process is analyzed separately.

To that end, we reprogrammed MEFs to iPSCs by OSKM and to iTSCs by GETM and examined their transcriptome (i.e. Bulk RNA-seq and SC-RNA-seq), methylome (i.e. RRBS), chromatin accessibility and activity (i.e. ATAC-seq and ChIP-seq for H3K4me2 and H3K27ac) and genomic stability (i.e. CNVs) at various time points along the process.

Initially, we asked whether the reprogramming process toward pluripotent and TSC states follows the same dynamics as of the forming cells during early embryogenesis. While clear transcriptional changes are found between the different stages (i.e. zygote, 2-cell stage, 4-cell stage, 8-cell stage, morula and blastocyst), the transcriptional heterogeneity within the cells of each group before blastocyst formation is relatively mild. This suggests a ‘T’-shaped model, where cells at each stage undergo relatively similar epigenetic and transcriptional changes before segregation, and dispersed into two distinct cells types, the ICM and TE, only at the morula/early blastocyst stage.

Our comparative and parallel multi-layer analysis revealed that, in contrast to cells during early embryogenesis that mostly resemble each other in each stage, cells undergoing reprogramming to pluripotent and TSC states exhibit unique and specific trajectories from the beginning of the process till the end, suggesting ‘V’-like behavior. Although similar processes such as somatic identity loss, proliferation, MET and metabolic shift occur in the two systems, each of the processes mostly uses different sets of genes and regulatory elements to induce its own fate. This ‘V’-shaped behavior was observed at all levels, starting from transcription and chromatin accessibility and activity and ending with DNA methylation.

We show that each reprogramming process uses different genomic regions and various strategies to silence fibroblastic identity. While the OSKM combination is very potent in inducing identity loss by interacting, from the onset of the process, with key regions that safeguard fibroblastic identity (i.e. regions that are enriched with ATF/CREB/AP1 sites), GETM open regions that are enriched with ATF/CREB/AP1 binding sites, which counteract their ability to silence the fibroblastic identity. This is in agreement with the SC-RNA-seq data that demonstrate a big fraction of cells with MEF-like identity even at day 12 of the reprogramming process by GETM.

By exploiting single-cell analysis, we demonstrate two unique and distinct populations of reprogrammable cells, suggesting that neither of the reprogrammable cells harbor a transcriptional profile that is shared during GETM and OSKM reprogramming. Moreover, we could also illuminate previously unknown reprogramming stages and markers for a faithful (Tdgf1 for OSKM and Cd82 for GETM) and failed (Anx3 for OSKM) reprogramming process for the two reprogramming systems.

These results clearly demonstrate that the reprogramming process of somatic cells toward pluripotency and TSC state takes completely different routes from the onset of reprogramming, and that somatic nuclear reprogramming and reprogramming during early embryonic development toward pluripotency and TE state are characterized by different properties and follow diverse paths.

However, we believe that key features that characterize the process of nuclear reprogramming by OSKM and GETM are shared with the reprogramming process that occurs before lineage specification in the early embryo. As such, by comparing OSKM to GETM reprogramming we could subtract all the general regions that are being remodeled in the fibroblastic nucleus during reprogramming at large and to focus on regions that are being remodeled specifically by OSKM factors. Remarkably, we identified a clear and significant embryonic development program that is executed by OSKM and involves chromatin remodeling of two of the most important organs of the developing embryo, the brain and the heart. Initially, OSKM demethylate, define and open these regions and subsequently limits their activity by decreasing the levels of the active histone mark, H3K27ac. In contrast, GETM factors induce DNA methylation on key developmental genes and thus shutting off this early embryonic development program. By inducing chromatin accessibility and activity and by increasing the levels of the active histone mark H3K27ac, GETM activate the trophoblastic program that involves the activation of metabolic processes that participate in transcription and translation, as well as migration and endothelial cell attraction, which are all known properties of trophoblast cells.

Overall, this study describes and illuminates key features that characterize the reprogramming process toward pluripotent and TSC states at all levels of regulation (i.e. DNA methylation, chromatin accessibility and activity, transcriptome and CNVs). By comparing two reprogramming processes simultaneously we were able to reveal new properties for the induction of pluripotent and TSC states, elements which could not have been elucidated had each reprogramming process been analyzed separately. We believe that the generation of such a database is a powerful tool to study cellular plasticity and cell fate decision.

**Table 1.**
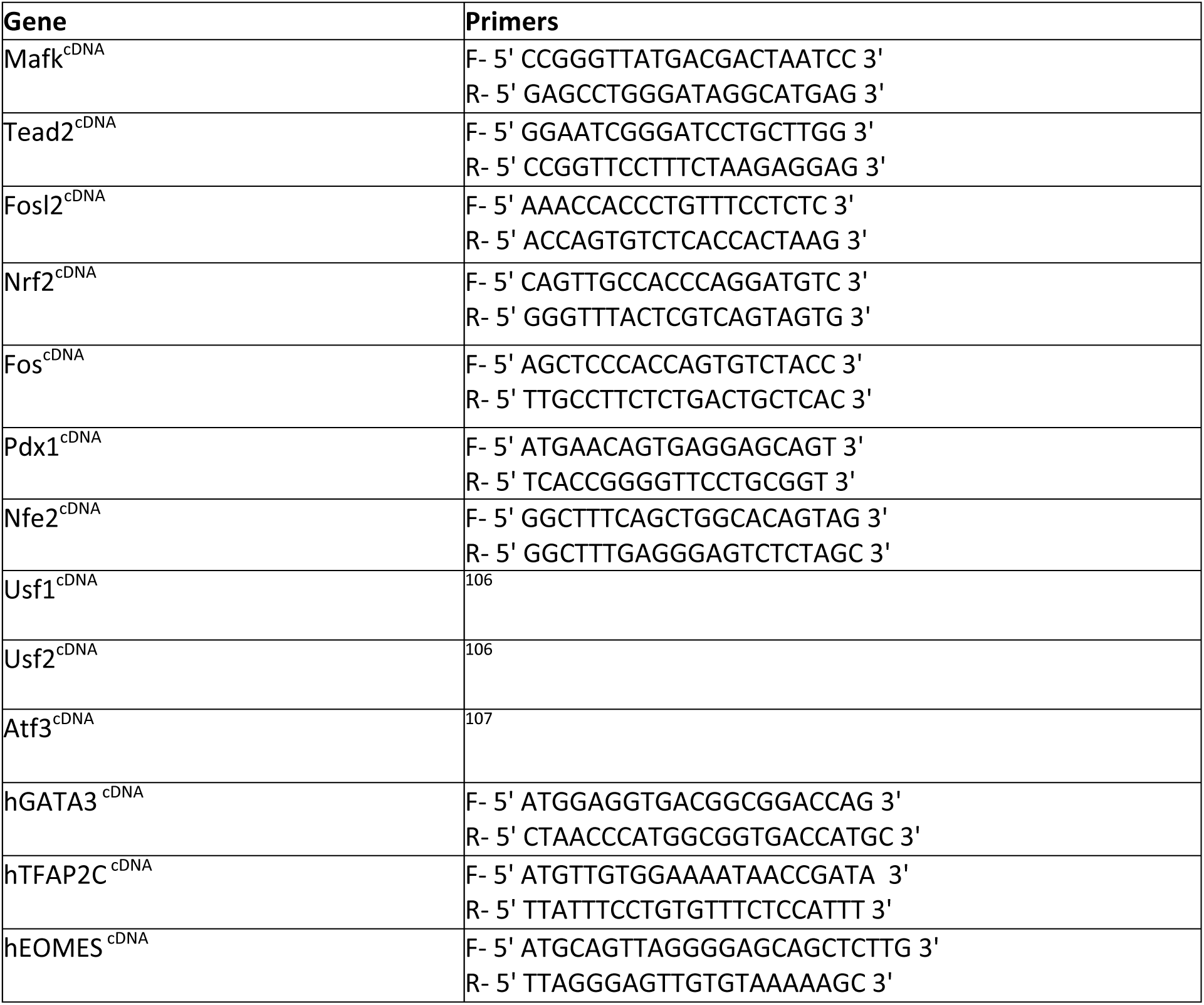
List of primers

## Acknowledgment

Y.B. is supported by research grants from the European Research Council (ERC, #676843), the Israel Science Foundation (ISF, #823/14), EMBO Young Investigator Programme (YIP), DKFZ-MOST (CA 177), Howard Hughes Medical Institute International Research Scholar (HHMI, #55008727) and by a generous gift from Ms. Nadia Guth Biasini. T.K. is supported by the Israel Science Foundation (ISF, #913/15 and #1250/18). A.R. is supported by the British GROWTH fellowship. N.L. is supported by CIDR Data Science and Leibniz fellowships

## Author Contribution

Y.B. conceived the study, prepared the figures and wrote the manuscript. Y.B. and M.J. designed the experiments. M.J. performed all the reprogramming experiments toward iTSCs and iPSCs, collected and prepared cells for all analyses and performed ChIP-seq, RRBS and cloned the various factors for reprogramming efficiency experiments. A.R. performed the bioinformatics analyses including bulk RNA, SC-RNA-seq, RRBS (together with N.L. and T.K.), ATAC-seq and ChIP-seq (together with N.L. and T.K.).

M.A. performed the reprogramming efficiency experiments with the various identified factors and scored and counted the colonies. S.S. prepared all the libraries for the RNA-seq and ATAC-seq and performed the reprogramming experiments for CNV analysis. T.K., M.Z. and A.E. performed and analyzed the CNV experiments on reprogrammable cells and fully reprogrammed iPSC colonies. T.K. supervised A.R. and N.L. and contributed to the design of the analyses and performed data integration analysis (together with A.R.). Y.B., A.R. and T.K. contributed to the interpretation of the results. K.M. injected BYKE cells to blastocysts for the purpose of MEF isolation.

## Author Information

All ATAC-seq, ChIP-seq, RRBS, SC-RNA-seq and bulk RNA-seq data is currently being deposited in the Gene Expression Omnibus database (GEO). We will be happy to share the data with the reviewers. Please let us know if you request it for the review process. Correspondence and requests for materials should be addressed to Y.B.

## Methods

### Cell culture and primary MEFs production

All ESCs and iPSCs were cultured in mouse embryonic stem cell medium containing 500ml DMEM supplemented with (15% FBS, 2mM L-Glutamine, 1% non-essential amino acid, in-house mouse Leukemia inhibitory factor (mLif), 0.1mM β-mercaptoethanol (Sigma), 1% penicillin-streptomycin and with or without 2i-PD0325901 (1 μM) and CHIR99021 (3 μM) (PeproTech). All TSCs and iTSCs were grown in TSC medium containing a combination of 70% MEF conditioned medium and 30% freshly prepared medium, (RPMI supplemented with 20% FBS, 0.1 mM β-mercaptoethanol, 2 mM L-Glutamine, 1% penicillin-streptomycin, 25 ng/ml human recombinant FGF4 (PeproTech) and 1 μg/ml heparin (Sigma-Aldrich).

Mouse embryonic fibroblasts (MEFs) were isolated as previously described^87^. Briefly, Pluripotent mouse ESCs and iPSCs were injected into E3.5 blastocysts, chimeric embryos were isolated at E13.5 and then dissected under the binocular to remove any internal organs and heads. The tissue was chopped by scalpels and incubated 30 minutes with 1ml Trypsin-EDTA (0.25%, Gibco) at 37°C. Next, trypsin activation was neutralized by 10ml DMEM containing 10% serum and the chopped embryos underwent intensive pipetting until homogeneous mixture of cells was noted. Each embryo was seeded into one 15cm plate and cultured with DMEM containing 10%FBS, 1% penicillin-streptomycin and 2mM L-glutamine. The cells were grown till the plate being full. Puromycin (2µg/ml) was added for selection for BYKE MEFs (the M2rtTA cassette that resides inside the rosa26 locus of the injected cells contains a resistance gene for puromycin), killing only the host cells. All cells were maintained in a humidified incubator at 37°C and 6% CO2.

### Molecular Cloning, Lentiviral Infection, and Reprogramming

The open reading frame of the examined genes (i.e. Ctcf^23^, Cdx2^13^, Atf3^88^, Tead2, Fosl2, Pdx1, Nrf2, Usf1, Usf2, NE-F2, Fos, MafK) was cloned into pMINI vector (NEB) and then restricted with EcoRI or MfeI and transferred into FUW-TetO expression vector. Lentiviruses were generated by transfecting vector DNA, (hGETM 3:3:3:1) or STEMCCA cassette for hOSKM, with a mix of lentiviral packaging vectors (7.5 µg psPAX2 and 2.5 µg pGDM.2) into 293T cells, the viruses were collected at 48, 60 and 72h after transfection, the medium containing the viruses was supplemented with 8 µg/ml of polybrene (Sigma) and filtered by 0.45 µm filter, the viruses were then added to MEFs (passage 0) that were seeded at 70% confluency two days prior to the first infection. Six hours following the third infection, medium was changed into DMEM containing 10% FBS. Eighteen hours later, medium was changed into reprogramming medium; ESC reprogramming medium (DMEM supplemented with 10%FBS, 0.1mM β-mercaptoethanol, 2mM L-glutamine, 1%non-essential amino acids, in-house mouse Leukemia inhibitory factor (mLif), and 2 μg/ml doxycycline) or TSC reprograming medium (RPMI supplemented with 20%FBS, 0.1mM β-mercaptoethanol, 2mM L-glutamine, in house mouse recombinant FGF4 (equivalent to 25ng/ml), 1 μg/ml heparin (Sigma-Aldrich), and 2μg/ml doxycycline). The two reprogramming mediums were changed every other day. For iPSC reprogramming, the MEFs were exposed to doxycycline for 15 days, followed by 5 days of dox withdrawal in ESC culturing medium. For iTSC reprogramming, the MEFs were exposed to doxycycline for 20 days, followed by 10 days of dox removal in TSC culturing medium. iTSCs colonies were then isolated, trypsinized, and plated in a well in a 6-well plate on feeder cells and passaged until stable colonies emerged.

### FACS analysis

Cells were trypsinized, washed with PBSx1 and filtered through mesh paper. Flow cytometry analysis was performed on a Beckman Coulter and cell sorting was performed on FACS-Aria III.

### RNA libraries and sequencing

Total RNA was isolated using the Qiagen RNeasy kit. mRNA libraries were prepared using the SENSE mRNA-seq library prep kit V2 (Lexogen), and pooled libraries were sequenced on an Illumina NextSeq 500 platform to generate 75-bp single-end reads.

### Reduced Representation Bisulfite sequencing (RRBS)

RRBS assay was performed as previously described^89^, briefly, 20ng of genomic DNA were digested with Msp1 restriction enzyme (NEB, R0106L), DNA fragments were end-repaired and A-Tailed using Klenow fragment (3’-5-exo-) (NEB, M0212L), the DNA fragments were ligated to illumina adaptors (Illumina, PE-940-2001) using T4 ligase (NEB, M0202M) and then size selected using AMPure XP beads (Beckman Coulter Genomics, A63881) The samples were then subjected to two consecutive bisulfite conversions using EpiTect Bisulfite Kit (QIAGEN, 59104) and PCR using PfuTurbo Cx hotstart DNA polymerase (Agilent Technologies, 600412). The RRBS libraries were sequenced by Illumina HiSeq 2000 platform.

### Single-cell RNA seq

Reprogrammable cells at day 6 or 12 were prepared as instructed in the 10X Genomics cell preparation guidelines. Briefly, cells were trypsinized and centrifuged at 1000 RPM for 3 minutes, then were washed twice with PBSx1 containing 0.04% BSA and cleaned from cell debris and large clumps by filtering throw mesh paper. Next, resuspended cells were subjected to dead cell removal kit (MACS, 130-090-101) to remove any non-viable cells. Cell viability were estimated using trypan blue staining. Cells were then resuspended in PBSx1 with 0.04% BSA at the concentration 1000 cells/µl and 4000 cells from each condition were subjected to 10x Genomics. Single-cell RNA libraries were prepared using Chromium Single Cell 3’ Library Kit v2 (10X genomics, 120234) and the generated libraries were sequenced using Illumina NextSeq 500 platform.

### Chromatin immunoprecipitation (ChIP)

Chromatin immunoprecipitation (ChIP) assay was performed as previously described^90^. Briefly, cells were fixed for 10 min at RT with a final concentration of 0.8% formaldehyde. Formaldehyde was quenched with glycine for a final concentration of 125mM. The cells were then lysate with lysis buffer (100mM Tris-HCl, 300mM NaCl, 2% Triton® X-100, 0.2%v sodium deoxycholate, 10mM Cacl2) supplemented with EDTA free protease inhibitor Roche-11873580001 for 20 min at Ice and the chromatin was digested by MNase (micrococcal nuclease)-Thermo Scientific™-88216 for 20 min at 37°C. MNase was inactivated by 20mM EGTA. The fragmented chromatin was incubated with pre-bounded Dynabeads (A and G mix) -Invitrogen 10004D/ 10002D using H3K27ac antibody (Abcam, ab4729) and H3K4me2 antibody (Millipore, 07-030). Samples were then washed twice with RIPA buffer, twice with RIPA high salt buffer (NaCl 360mM), twice with LiCl wash buffer (10mM Tris-Hcl, 250mM LiCl, 0.5% DOC, 1mM EDTA, 0.5% IGEPAL), twice with 10mM Tris-HCl pH=8. DNA was purified by incubating the samples with RNAse A (Thermo Scientific™ EN0531) for 30 min at 37°C followed by a 2 hours incubation with Proteinase K (Invitrogen™ 25530049). DNA was eluted by adding 2X concentrated elution buffer (10mM Tris-HCl, 300mM NaCl, 1% SDS, 2mM EDTA) and then reverse crosslinked overnight at 65°C. Finally, DNA was extracted using AMPure XP beads (Beckman Coulter Genomics, A63881). Chip sample libraries were prepared according to Illumina Genomic DNA protocol as described^91^.

### ATAC libraries and sequencing

ATAC-seq library preparation was performed as previously described^92^. Briefly, cells were trypsinized and 50,000 cells were counted and incubated in lysis buffer to isolate nuclei. Nuclei were then resuspended in transposase reaction mix for 30 min at 37 °C (Illumina, Fc-121-1030). The samples were purified using Qiagen MiniElute kit (QIAGEN, 28204), Transposed fragments were directly PCR amplified and sequenced on an Illumina NextSeq 500 platform to generate 2 × 36-bp paired-end reads.

### Data processing

#### A) Bulk RNA-seq

Low quality bases and sequencing adaptors of 36 raw fastq files RNA-seq containing single-end 61bp-long reads were trimmed using Trim Galore (V 0.6.0, https://github.com/FelixKrueger/TrimGalore) and then mapped to the mm9 reference genome using HISAT2 (V 2.1.0, ^93^) with default parameters. Read counting was performed using featureCounts (V 1.6.2, ^94^) with (Mus_musculus.NCBI37.gtf annotation). Differential gene expression analysis was performed using DESeq2_1.26.0 package^94^. Unsupervised hierarchical clustering was performed for 10,000 most variable genes among ESCs, bdTSCs, fibroblasts and cells during reprogramming. R package dynamicTreeCut ^95^ was used to perform adaptive branch pruning detecting 27 prominent clusters. R packages Enrichr (V 2.1, ^96^) and ClusterProfiler (V 3.14.3, ^97^) were used to query Biological processes, Mouse gene atlas and KEGG pathways analysis of significantly over-represented genes for each cluster. A second aligner TopHat [4] (v2.0.6, ^98^) was used to map reads to mm9 reference genome. Mapped reads were then processed using cufflinks [4] (v2.0.2, ^99^), and gene expression levels (FPKM) were calculated for each replicate.

#### B) 10x Single-cell data

SC-RNA-seq libraries were generated from each time point using the 10X Genomics. The cellranger-3.0.2, https://github.com/10XGenomics/cellranger was used for mapping of the 10x single-cell RNA-seq data. Read1 data of pooled cells were split into single-cell data using the barcode sequences contained in the first 16 bps. The next 10 bps were recorded as unique molecular identifiers (UMIs). Read2 with 75 bp were aligned to the mm10 reference genome. We used Seurat (V 3.1.4, ^100^) to pre-processing the data and perform clustering. The function ‘FindAllMarkers’ to identify the marker genes for each of the clusters in the UMAP representation. For day 6 OSKM and GETM reprogramming, we excluded cells with detected genes less than 2000 or cells with sum of the non-normalized UMI counts less than 10,000, or cells with percentage of mitochondrion UMI values larger than 10%. For day 12 OSKM and GETM reprogramming, we excluded cells with detected genes less than 1000 or cells with sum of the non-normalized UMI counts less than 5000, or cells with percentage of mitochondrion UMI values larger than 15%. R package DoubletFinder (https://github.com/chris-mcginnis-ucsf/DoubletFinder) was used to identify and exclude potential doublets. Overall, 5899 cells of day 6 OSKM-GETM reprogramming and 5752 cells of day 12 OSKM-GETM passed the quality control criteria. On median, there were 16232/13488 UMI counts and 4079/3735 detected genes for each cell of day6/day12 OSKM-GETM dataset.

### DNA methylation

Low quality bases and sequencing adaptors of 45 raw fastq files were trimmed using Trim galore (V 0.6.0, https://github.com/FelixKrueger/TrimGalore) and then mapped to the mm9 reference genome using Bsmap (V 2.90, ^100^) with flags -S 10 -R -p 8 -D C-CGG. Bam files belonging to same reprogramming system and day were merged to ensure maximum overlap between all samples. Methylation beta values were extracted from the BAM files using wgbs_tools (https://github.com/nloyfer/wgbs_tools). Methylation markers were identified using in-house developed script find_markers.py to generate Bed files with p-value < 0.05 between different conditions summarized in different groups. 130,000 blocks were identified with significant methylation alteration that occurs during reprogramming in both OSKM and GETM reprogramming. In order to minimize noise and extract significant trends, we used the K-means algorithm to classify ∼130,000 blocks that are shared amongst all samples during reprogramming to a TSC or pluripotent states and obtained 100 clusters. A new table was constructed by averaging DNA methylation levels per sample per cluster and then projected the processed data onto the first two principal components. Clusters loading plot showed significant clusters contributed to the first two principal components and clusters that are near to each other showed similar trends of methylation allowing us to extract 15 different trends shown as heatmaps in Fig. 4A and Fig. S4A. Genomic regions associated with all blocks belonging to each of the 15 clusters were annotated using GREAT (V 4.0.4^48^) and were summarized in Supplementary Table 2.

### ATAC-seq

Fastq files were mapped to the mm9 reference using bwa (https://arxiv.org/abs/1303.3997, version 0.7.17-r1188). The mapped reads were converted to BAM format and filtered by mapping quality (MAPQ) of >=10, retaining only properly aligned pairs (samtools -F 1796 flag). The BAM files were then sorted and indexed using samtools (v1.9 ^101^).

Bigwig coverage tracks were generated using deepTools bamCoverage (v3.4.1 ^102^) with the following flags: --normalizeUsing RPGC -bs 50 -e 500 --effectiveGenomeSize 2150570000.

Coverage peaks were called using MACS (v2.1.2 ^103^) with flags -g mm --slocal=2000 --llocal=20000 -- nomodel --extsize=300 -f BAMPE.

Peaks of multiple replicates were retained only if identified by 30% of the replicates, or more.

### ChIP-seq

Fastq, BAM and bigwig files were processed in a similar way to the ATAC-seq files.

### Annotation of genomic regions

Peaks from each experiment were then divided into subsets, including peaks that appear in both OSKM and GETM (3, 6, or 9 days after induction) but not in MEFs, peaks from GETM (days 3,6,9) not identifiable in MEFs, OSKM peaks (days 3, 6, 9) not identifiable in MEFs, and disjoint sets of cell-type specific peaks (e.g. GETM day 3 peaks not found in MEFs or in OSKM day 3, etc.). We also analyzed peaks from ESC, TSC or MEF cells.

Genomic regions from each group of peaks were then annotated using annotatePeaks.pl (HOMER suite, http://homer.ucsd.edu/homer/ngs/annotation.html, UCSC mm9 genome version) as Promoter, TTSs, 5’ and 3’ UTRs, or as Exonic, Intronic, or Intergenic regions.

### Motif analysis

ATAC-seq peaks were called in each replicate separately, and overlapping peaks from replicates were then merged. Peaks overlapping MEF peaks (top 50K) were then removed. Finally, the center 250bp of each peak was considered for further analysis (peaks shorter than 250bp were removed).

We them further divided the peaks of each time point into disjoint groups, including peaks identified in both GETM and OSKM ATAC-seq (e.g. GETM&OSKM D03), GETM-only peaks (e.g. GETM\OSKM D03) or OSKM-only peaks (e.g. OSKM\GETM D03).

Finally, we used “findMotifsGenome.pl -nomotif” (HOMER suite) to identify occurrences of known motifs within those sequences. A similar approach was applied to H3K27ac and H3K4me2 ChIP-seq peaks.

### CNV analysis

Read alignment was done with BWA mem 0.7.15 ^104^ to the mouse reference genome including PhiX174. Copy number analysis was performed using library cn.mops^105^ in paired mode for 4kb windows and applying DNAcopy for segmentation with R (R Core Team (2015). R: A language and environment for statistical computing. R Foundation for Statistical Computing, Vienna, Austria. https://www.R-project.org/).

## SUPPLEMENTARY

**Supplementary Fig. 1.**
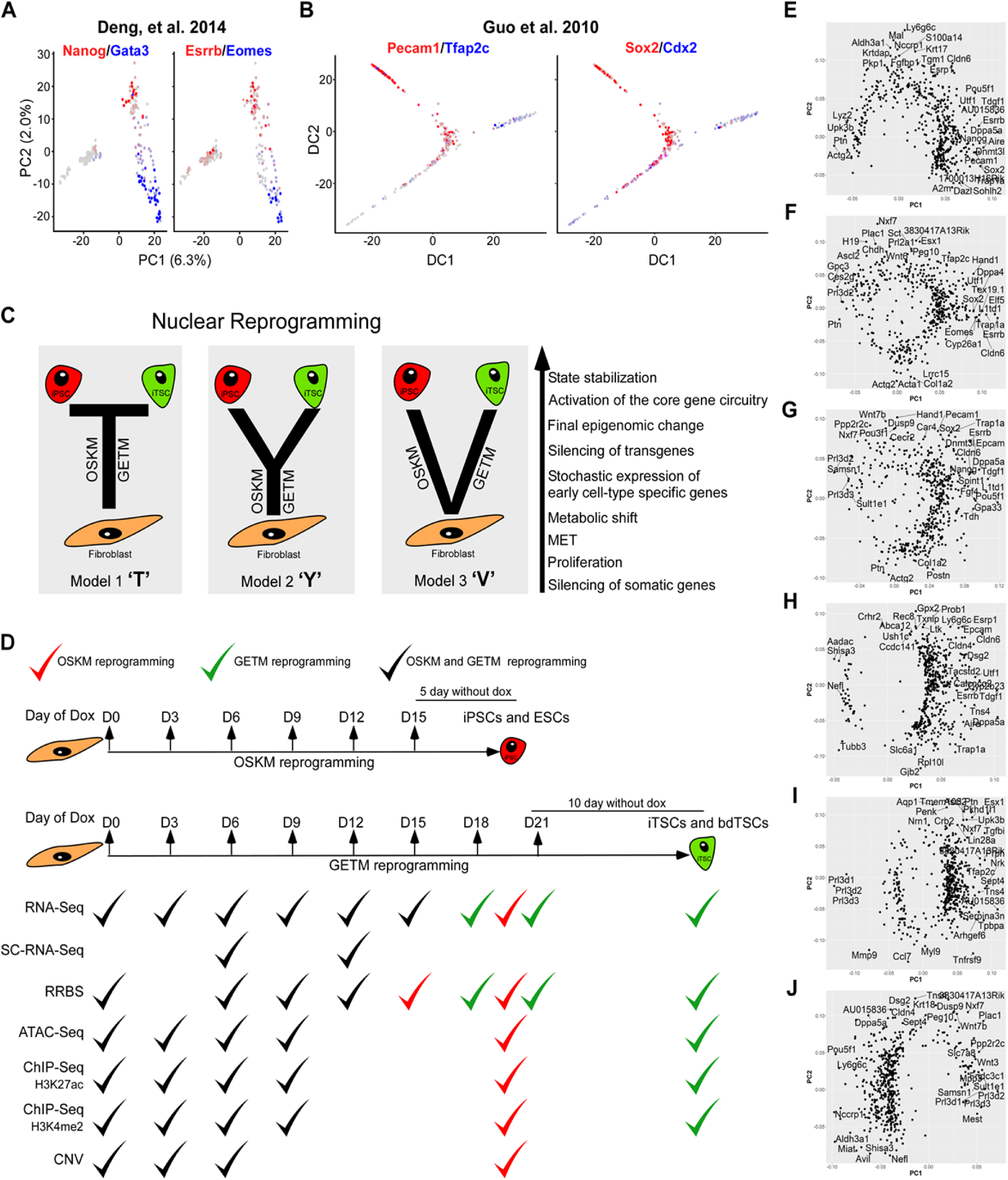
Embryonic development trajectory and hypothetical nuclear reprogramming progression models using comparative multi-omics analysis. **(A)** Overplayed co-expression of Nanog/Gata3 and Esrrb/Eomes among individual cells extracted from^5^. Red color indicates cells that are enriched with key pluripotency genes while blue color indicates cells with enrichment of key TE genes. **(B)** Overplayed co-expression of Pecam1/Tfap2c and Sox2/Cdx2 among individual cells extracted from^7^. Red color indicates cells that are enriched with key pluripotency genes while blue color indicates cells with enrichment of key TE genes. **(C)** Schematic illustration of three possible models, ‘T’, ‘Y’, ‘V’, explaining the reprogramming progression of fibroblasts toward either iPSCs by OSKM or iTSCs by GETM. **(D)** Schematic representation of the reprogramming process of fibroblasts to iPSCs (top, red) and iTSCs (bottom, green) and the various high throughput experiments and time points that were analyzed. Black ‘V’ represents a time point that was taken for both GETM and OSKM reprogramming while Green ‘V’ represents GETM-only time point and red ‘V’ represents OSKM-only time point. **(E-G)** PCA loading plots showing the contribution of individual genes out of top 500 most differentially expressed genes to the first and second PCA components; distance from the origin along each axis corresponds to strength of contribution to that component. Shown are PCA loading plots for the reprogramming process to either iPSCs (E), iTSCs (F) or both (G) as assessed by gene expression profiles of bulk RNA-seq data projected onto the first two principal components. 32 highest ranked genes are marked in each plot. **(H-J)** same as in (E-G) but here only reprogrammable cells (cells on dox) are plotted.

**Supplementary Fig. 2.**
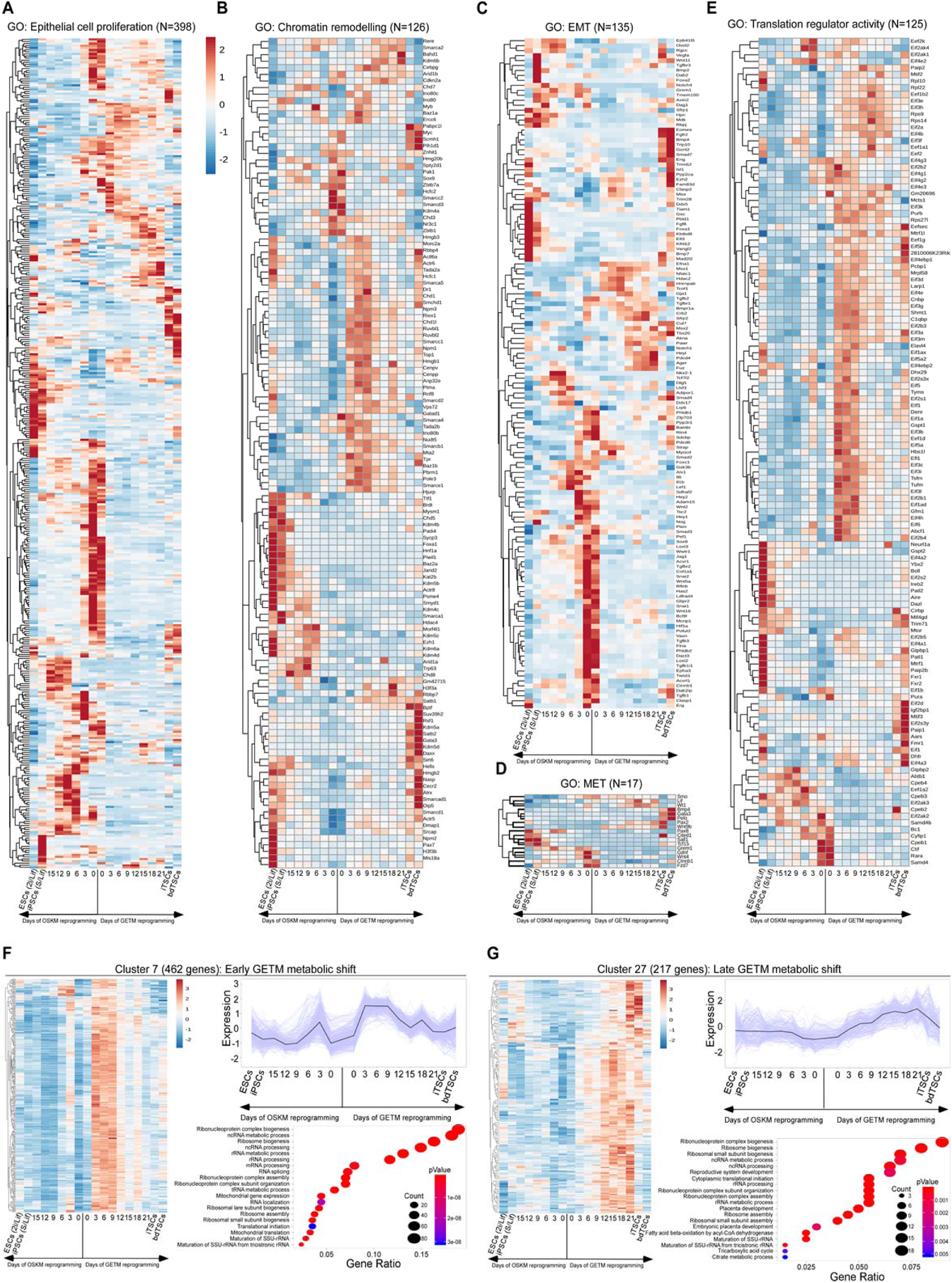
GETM and OSKM reprogramming factors mostly exhibit mutually exclusive transcriptional profiles during reprogramming. **(A-E)** Heatmaps showing the expression levels of genes involved in early and general reprogramming processes such as epithelial cell proliferation (A), chromatin remodeling (B), EMT (C), MET (D) and translation regulator activity (E), during the conversion of fibroblasts into iPSCs by OSKM and into iTSCs by GETM. **(F)** Heatmap, expression pattern plot and GO terms of 462 genes of cluster #7 as detected by bulk RNA-seq during reprogramming toward iPSCs and iTSCs. **(G)** Heatmap, expression pattern plot and GO terms of 217 genes of cluster #27 as detected by bulk RNA-seq during reprogramming toward iPSCs and iTSCs.

**Supplementary Fig. 3.**
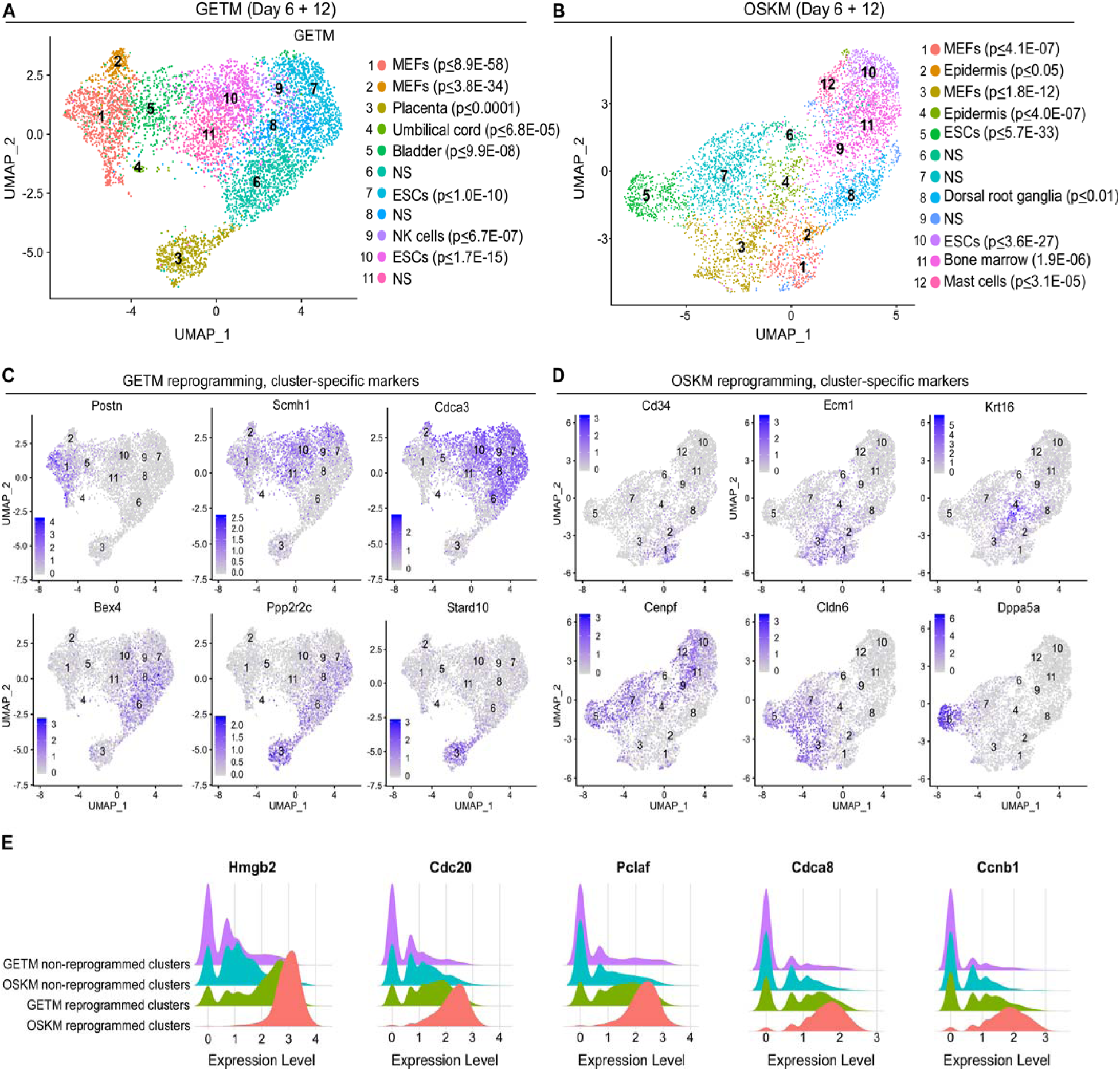
Single-cell RNA-seq analysis on OSKM and GETM reprogrammable cells identifies reprogramming and stage-specific markers. **(A)** UMAP visualization analysis of both 3,288 single cells at day 6 and 3,029 single cells at day 12 of GETM reprogramming. Marker genes were used to characterize all subpopulation and using the mouse gene atlas only the closest significant cell type was assigned to each subpopulation. NS refers to non-significant. **(B)** UMAP visualization analysis of both 2,611 single cells at day 6 and 2,723 single cells at day 12 of OSKM reprogramming. Marker genes were used to characterize all subpopulation and using the mouse gene atlas only the closest significant cell type was assigned to each subpopulation. **(C-D)** Expression level of selected cluster-specific markers during GETM reprogramming (days 6 and 12) and OSKM reprogramming (days 6 and 12), respectively. The expression level of the specified markers is visualized by a range of intensities of a purple color. **(E)** Redge plots showing the expression distributions of key proliferation genes where each community represents potentially reprogrammed or non-reprogrammed cells for each reprogramming system.

**Supplementary Fig. 4.**
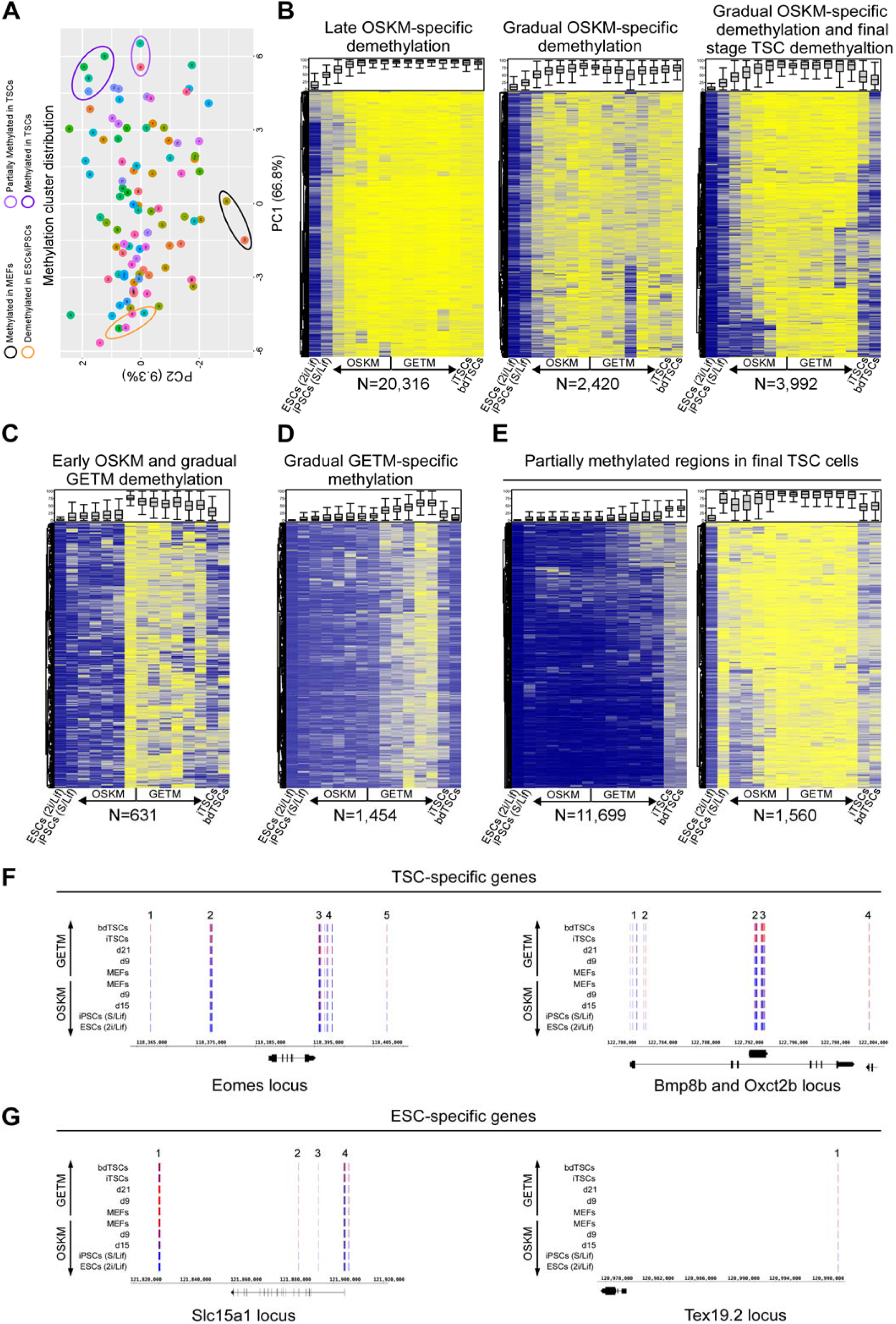
Methylation dynamics during OSKM and GETM reprogramming. **(A)** PCA plot projected by the first two principal components showing the average DNA methylation per sample for 100 clusters obtained using K-Means clustering algorithm of 130,000 blocks. Clusters that are near to each other show similar trend of methylation. **(B-E)** Heatmaps showing the dynamics of DNA methylation alterations and specific cluster patterns across bulk samples during reprogramming towards both pluripotent and TSC states. **(F)** DNA methylation level of genomic loci that contain gene specific to TSCs (e.g. Eomes, Bmp8b and Oxct2b). **(G)** DNA methylation level of genomic loci that contain gene specific to ESCs (e.g. Slc15a1 and Tex19.2).

**Supplementary Fig. 5.**
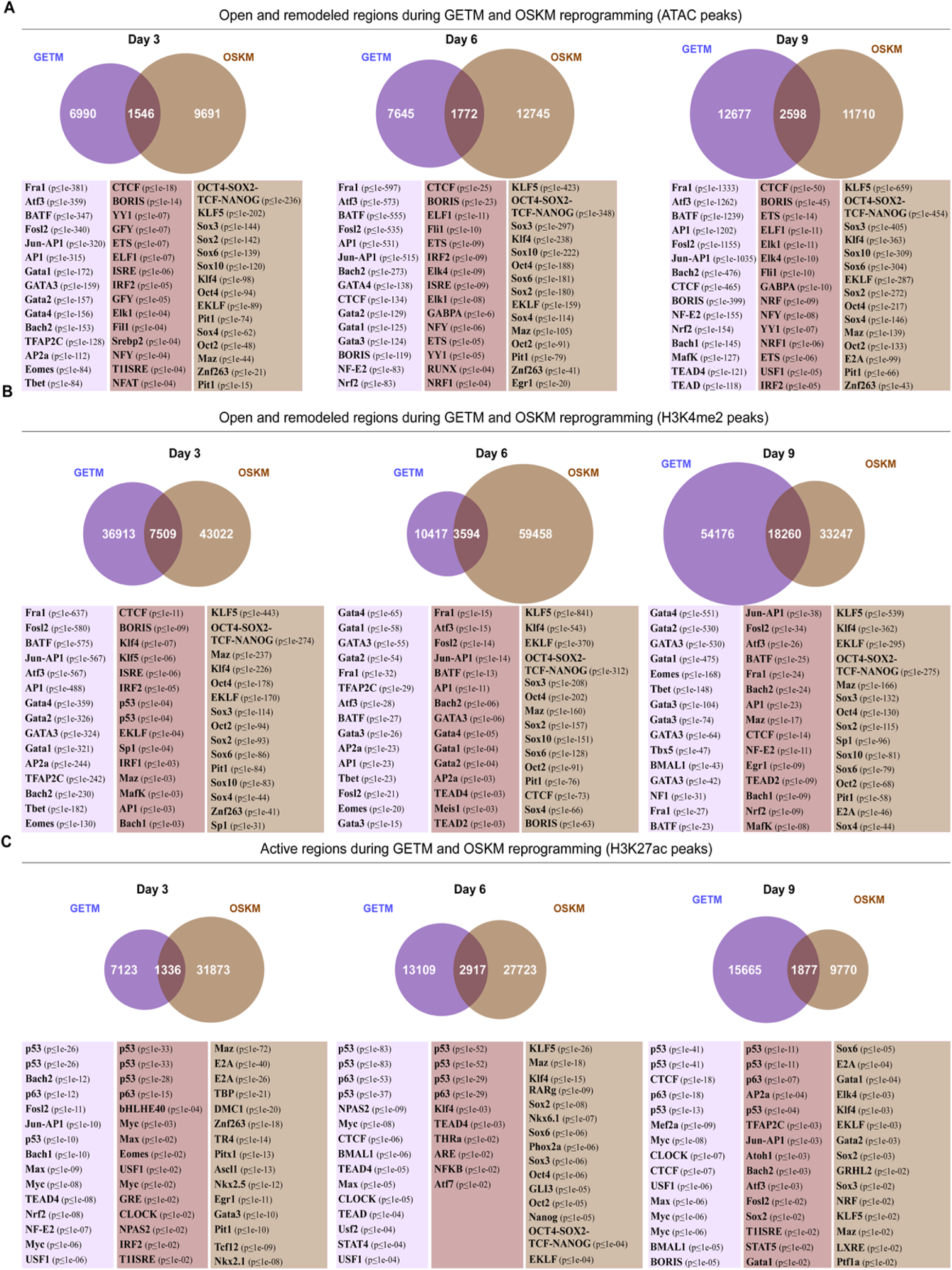
Motif enrichment of ATAC, H3K27ac and H3K4me2 peaks during GETM and OSKM reprogramming. **(A-C)** Venn diagrams and motif analysis for ATAC-seq peaks (A), H3K4me2 peaks **(A)** and H3K27ac peaks (C). Comparison of GETM-only (left wedge, purple), GETM and OSKM (interaction), and OSKM-only (right wedge, brown) peaks from day 3 to day 9. Below are motifs, differentially enriched between each set of peaks vs. the rest.

**Supplementary Fig. 6.**
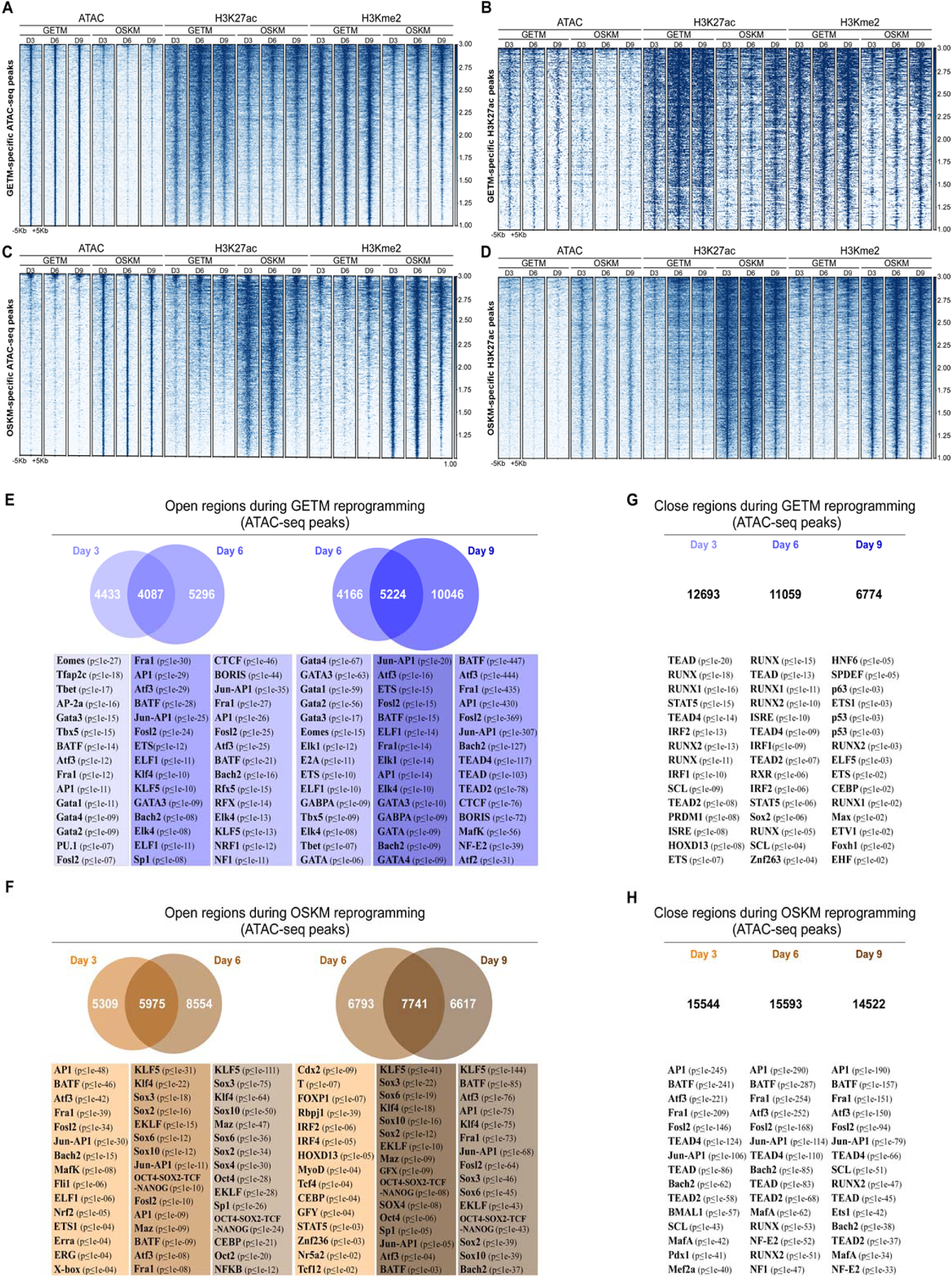
Unique behavior and motif enrichment of ATAC, H3K27ac and H3K4me2 peaks during GETM and OSKM reprogramming. **(A-D)** Heatmap showing ATAC-seq and ChIP-seq (H3K27ac and H3K4me2) across 1716 differential ATAC-seq peaks (p<1e-3) in GETM (A and B) or 2848 OSKM (C and D). Shown are genomic regions of peak locations ±5Kb. Differential peaks were called using DESeq2 analysis on the number of reads in each of 18,421 ATAC-seq peaks, using a significance threshold of adjusted p-value < 1e-3. **(E)** Comparison of GETM ATAC-seq peaks from days 3 and 6 (left) or 6 and 9 (right). Shown below are enriched motifs for each binary set of peaks. From left to right: day 3-only (left wedge) vs all day 6 peaks; day 3 and 6 peaks (intersection) vs day 3-only and day 6-only peaks; day 6-only (right wedge) vs all day 3 peaks; day 6-only (left wedge) vs all day 9 peaks; day 6 and 9 peaks (intersection) vs day 6-only and Day 9-only peaks; day 9-only (right wedge) vs all day 6 peaks. **(F)** Same as E but for OSKM **(G)** Similar analysis for 12,693 genomic regions that are accessible in MEFs but close on GETM day 3; or 11,059 regions that are accessible in day 3 but close in day 6; or 6774 regions that are accessible in Day 6 but close in day 9 **(H)** Same as G but for OSKM.

**Supplementary Fig. 7.**
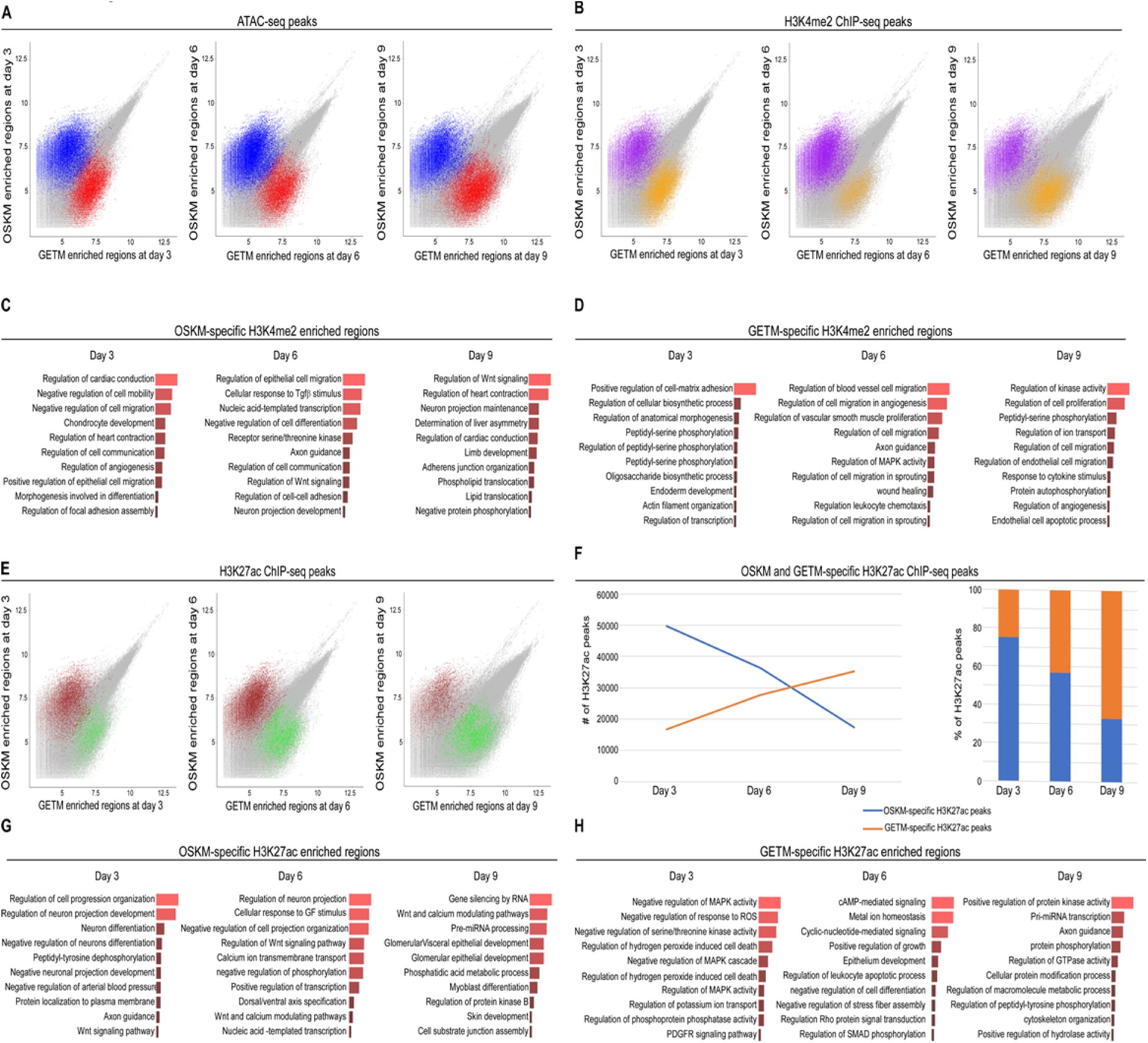
OSKM and GETM enriched regions for ATAC-seq peaks, H3K4me2 peaks and H3K27ac peaks. **(A)** Scatter plot of enriched regions differentially accessible between different cellular states during the reprogramming process using GETM and OSKM at days 3, 6, and 9. Differential accessible regions are marked with blue for OSKM, and red for GETM with adjusted p-value < 0.001. **(B)** Scatter plot of enriched regions that are both differentially accessible and enriched for H3K4me2 ChIP-seq differential peaks on top of the ATAC-seq signal during the reprogramming process using GETM and OSKM at days 3, 6 and 9. Enriched regions are marked with purple for OSKM and orange for GETM. **(C)** Top 10 enriched gene ontology (GO) terms within OSKM-specific regions that are both differentially accessible and enriched for H3K4me2 at days 3, 6 and 9 tested in the biological process ontology. **(D)** Top 10 enriched gene GO terms within GETM-specific regions that are both differentially accessible and enriched for H3K4me2 at days 3, 6 and 9 tested in the biological process ontology. **(E)** Scatter plot of enriched regions that are both differentially accessible and enriched for H3K27ac ChIP-seq differential peaks on top of the ATAC-seq signal for enriched regions during the reprogramming process using GETM and OSKM at days 3, 6 and 9. Enriched regions are marked with maroon for OSKM and green for GETM. **(F)** Line plot and stacked column chart showing differential dynamics of enrichment for H3K27ac during both OSKM and GETM reprogramming at days 3, 6, 9. **(G)** Top 10 enriched GO terms within OSKM-specific regions that are both differentially accessible and enriched for H3K27ac at days 3, 6 and 9 tested in the biological process ontology. **(H)** Top 10 enriched GO terms within GETM-specific regions that are both differentially accessible and enriched for H3K27ac at days 3, 6 and 9 tested in the biological process ontology.

**Supplementary Fig. 8.**
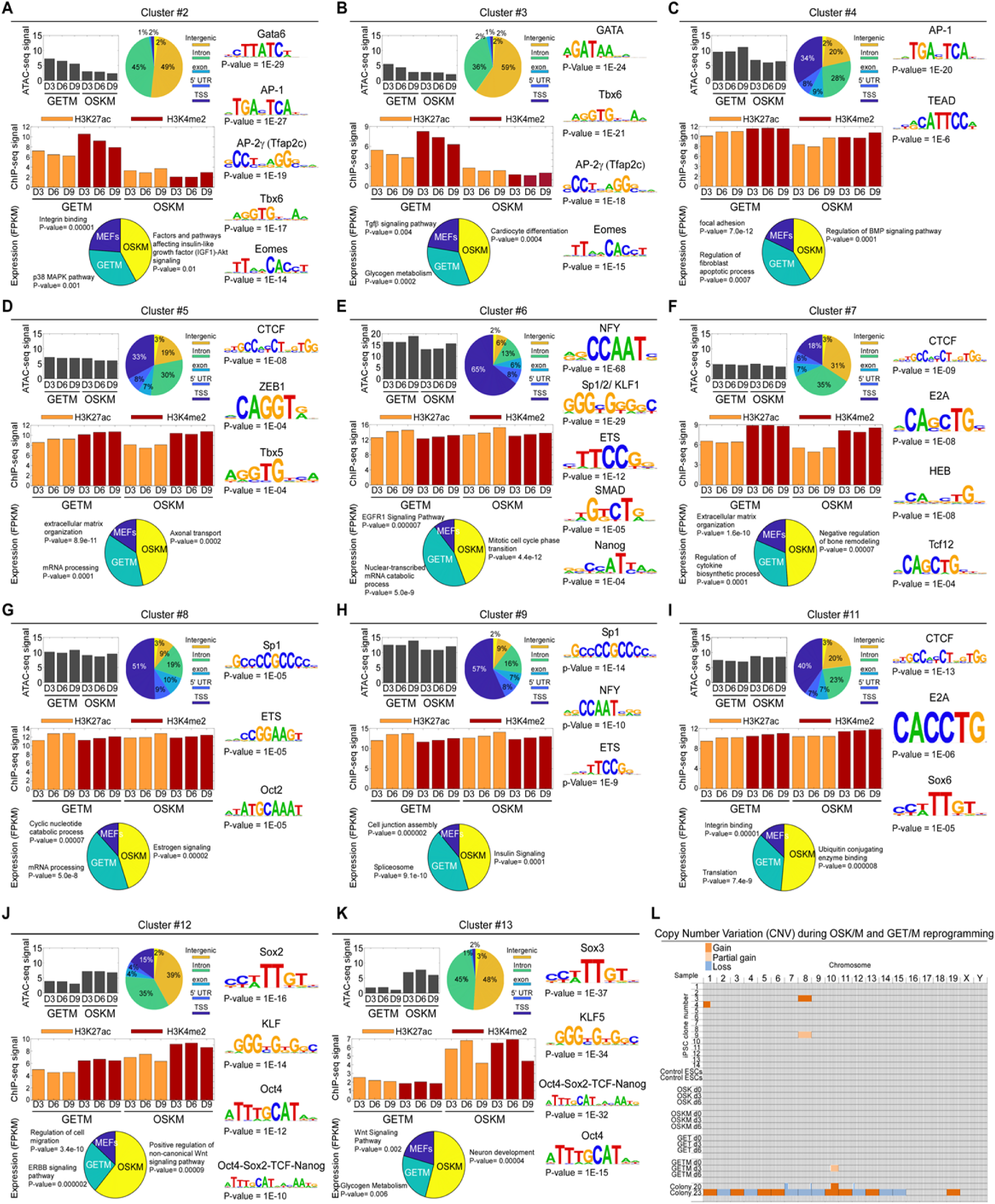
Data integration of chromatin accessibility and activity and gene expression and CNV analysis. **(A-K)** 18,420 GETM and OSKM ATAC peaks from days 3, 6, 9 were clustered to 14 clusters. Shown for each cluster are: mean ATAC-seq signal (top left), analysis of their genomic annotations (pie chart, center), enriched transcription factor motifs (right panel), average ChIP-seq signals of H3K27ac and H3K4me2 following GETM and OSKM induction (middle panel), and a pie chart for RNA expression levels and GO terms for genes that are associated with each cluster ATAC-seq peaks and exhibit the highest expression levels in MEFs (blue), or GETM (green) or OSKM (yellow, Bottom panel). **(L)** A graph summarizing the various copy number variations (CNVs) identified in OSK/M or GET/M reprogrammable cells (days 0, 3 and 6) and in isolated iPSC clones. Final TSCs/iTSCs hold an intrinsic capacity to accumulate genomic aberrations^13^ and thus are not measured here. Parental ESC line and partially reprogrammed iPSC colonies number 20 and 23^23^ were used as negative and positive control, respectively. All data were aligned to the parental MEFs.

